# The evolution of dependence and cohesion in incipient endosymbioses

**DOI:** 10.1101/2023.07.17.549359

**Authors:** Gaurav S. Athreya, Peter Czuppon, Chaitanya S Gokhale

## Abstract

Eukaryogenesis is the prototypical example of an egalitarian evolutionary transition in individuality, and endosymbiosis, more generally, is central to the origins of many complex biological systems. Why do only some symbioses undergo such a transition, and how does the host-symbiont relationship change during this process? Here, we characterise endosymbiosis by two emergent collective-level properties: host and symbiont survival as a collective (“mutual dependence”) and the level of synchronised reproduction (“reproductive cohesion”). Using adaptive dynamics, we study the evolution of the traits underlying these properties. First, by adding a carrying capacity for the collective population – a realism omitted in previous models – we find novel reasons why complete dependence or cohesion might not evolve, thus providing further theoretical support for the rarity of transitions in individuality. Second, our model suggests that asymmetries in evolutionary outcomes of hosts and symbionts can be explained by a difference in their population growth parameters, coupled with their shared fate when in a collective. Lastly, we show that during the early stages of an endosymbiosis, even if investments in dependence and cohesion are uncorrelated, mutual dependence arises faster than reproductive cohesion. Our results hence shed light on three aspects of endosymbiosis: coevolution between the host and symbiont, coevolution between dependence and cohesion, and ultimately on the opportunity of undergoing an evolutionary transition. Connecting to ecological factors, this work uncovers fundamental properties of endosymbioses, providing a clear way forward for theoretical and empirical investigations.

## Introduction

Endosymbiosis is a phenomenon of central importance in evolutionary biology, leading to the origin of eukaryotes and several astonishing long-term associations between different species. It is the prototypical example of an egalitarian transition in individuality: initially, autonomous and unrelated entities – the host and symbiont – come together to give rise to a more complex, integrated entity (1–4). The origin of endosymbiosis is not only a marked evolutionary transition but also a characteristic energy transition (5). Arguably, access to more energy processing via the endosymbionts set the stage for further evolutionary leaps such as multicellularity and beyond (6).

Endosymbionts are present in most life forms, even in unicellular prokaryotes (7, 8). Mitochondria and plastids have been famously shown to be endosymbionts arising from an ancient union between Archaeabacteria and an alphaproteobacterium (9–12). Many insects, such as the sap-sucking aphids, have been co-diversifying with their *Buchnera, Wigglesworthia*, and *Wolbachia* endosymbionts for millions of years (13– 16). There are examples from diverse ecologies across the globe - methanogenic endosymbionts in anaerobic ciliates (17), nitrogen-fixing endosymbionts in the diatom *Rhopalodia* (18), consortia of chemosynthetic bacteria in gutless tubeworms (19), and cyanobacterial endosymbionts in sponges (20). New kinds of endosymbiotic associations continue being discovered, such as denitrifying endosymbionts in anaerobic ciliates (21). Despite this widespread prevalence, endosymbiosis and egalitarian evolutionary transitions are understudied relative to their fraternal counterparts, such as the evolution of multicellularity or eusociality (for which there exist many review or book-length treatments, e.g. (22, 23)). However, this point of view is slowly changing with the recent emergence of more studies on symbiosis (24–29).

In this work, we are interested in the relationship between the host and symbiont and how it evolves throughout an evolutionary transition in individuality. By a transition in individuality, we mean the emergence – from individuals of one or more species that can undergo evolution – of a higher-level entity that can itself undergo Darwinian evolution. Transitions where the lower-level individuals making up a collective are unrelated are called “egalitarian” (30). Some symbioses have undergone such a transition, whereas others have not. For example, proto-mitochondria and their ancestral hosts underwent a transition to form the modern eukaryotic cell (9). *Buchnera* endosymbionts are obligately dependent on their aphid hosts, vertically transmitted, and have small genomes (31). On the other hand, while the tubeworm *Riftia* is obligately dependent on its symbiont *Endoriftia*, the latter has a free-living stage, and is transmitted horizontally (32). Similarly, the bobtail squid *Euprymna scolopes* and its bioluminescent *Vibrio* symbiont are both facultative, and the symbiont is acquired horizontally every generation (33). What controls this difference in outcomes, and how is it impacted by the host’s and symbiont’s life-history traits? How do the properties of the collective co-evolve, and is there a difference between host and symbiont evolutionary trajectories? Such questions were introduced in the works of (24) and (28). These models emphasized different aspects of symbioses: (24) considered the evolution of an exploitative symbiont and showed that even unidirectional resource transfer can lead to an evolutionary transition. (28) considered a tradeoff between independent reproduction and host encounter rate, and showed that this can lead to evolutionary equilibria where the symbiont is facultatively dependent on its host. However, certain assumptions were made for analytical tractability, such as exponential population growth without any regulation for hosts, symbionts, or collectives (24) and ignoring the evolution of host traits, and letting collectives grow exponentially (28). In this study, we, too, make a new simplifying assumption (see the Model section, parameter *d*), but it allows us to relax many others and, in doing so, ask a variety of different questions about host-symbiont co-evolution with reasonable population dynamics.

To begin, we define a host-symbiont collective to be an obligate endosymbiosis if it exhibits three properties: (i) intracellular location of the symbiont; (ii) at least one of the host or symbiont is obligately dependent on this interaction; and (iii) the collective can reproduce synchronously, i.e. as a unit. Following (34), we use “symbiosis” to mean any sustained organismal interaction on the pathogenic-beneficial continuum. Synchronised collective reproduction, which is the target of our notion of “reproductive cohesion”, is similar to, but stronger than, vertical symbiont transmission: we also include the requirement that the replication of the two partners is coupled, such as the coordination of mitochondrial fission and segregation with the cell cycle (although this is more complicated, see (35)). Importantly, synchronised reproduction endows the collective with a life cycle, previously proposed as the defining characteristic of an entity that can undergo an evolutionary transition (36). For example, our definition excludes gut microbiomes as there is no intracellular location or synchronised reproduction.

To study the conditions for the evolution of obligate endosymbiosis, we use evolutionary invasion analysis from the adaptive dynamics framework (37–40). This is a framework to study the long-term evolution by natural selection of traits that affect their bearer’s ecological interactions. The evolutionary fate of new rare mutants is studied by determining if they can invade and fix in the population in which they arise. This analysis assumes the separation of ecological and evolutionary timescales: the ecological processes dictating the fate of a mutant (“natural selection”) take place much faster than the timescale on which new host/symbiont mutants arise.

### An eco-evolutionary model

We consider three populations – free-living hosts (H), freeliving symbionts (S), and host-symbiont collectives (C). The primary process of interest is the evolution of the growth rates of these populations - we characterise obligate endosymbiosis by a positive growth rate for the collective and a zero growth rate for the free-living types.

Inspired by (25), we characterise a symbiosis by two emergent, collective-level quantities: the degree of host-symbiont mutual dependence and the degree of their reproductive cohesion (see Fig. 1). Mutual dependence is an aggregative measure of how well the two partners grow when free-living instead of when they are part of a collective; reproductive cohesion measures how often they reproduce synchronously instead of individually. The evolution of symbioses can thus be visualised as taking place in the plane of these two collective-level quantities. We say that an evolutionary transition in individuality occurs when there is complete mutual dependence and reproductive cohesion - the constituent individuals cannot live or reproduce without the other. We aim to understand the evolutionary trajectories in this dependence-cohesion plane. To study the joint evolution of the traits underlying dependence and cohesion, we use the method of evolutionary invasion analysis. The main object of interest is the invasion fitness of a mutant in the environment generated by a resident population. It is assumed in this framework that the ecological and evolutionary timescales can be separated, i.e., the (ecological) realisation of a mutant’s fate takes place much faster compared to the (evolutionary) timescale on which the next mutant arises (37).

**Fig. 1.**
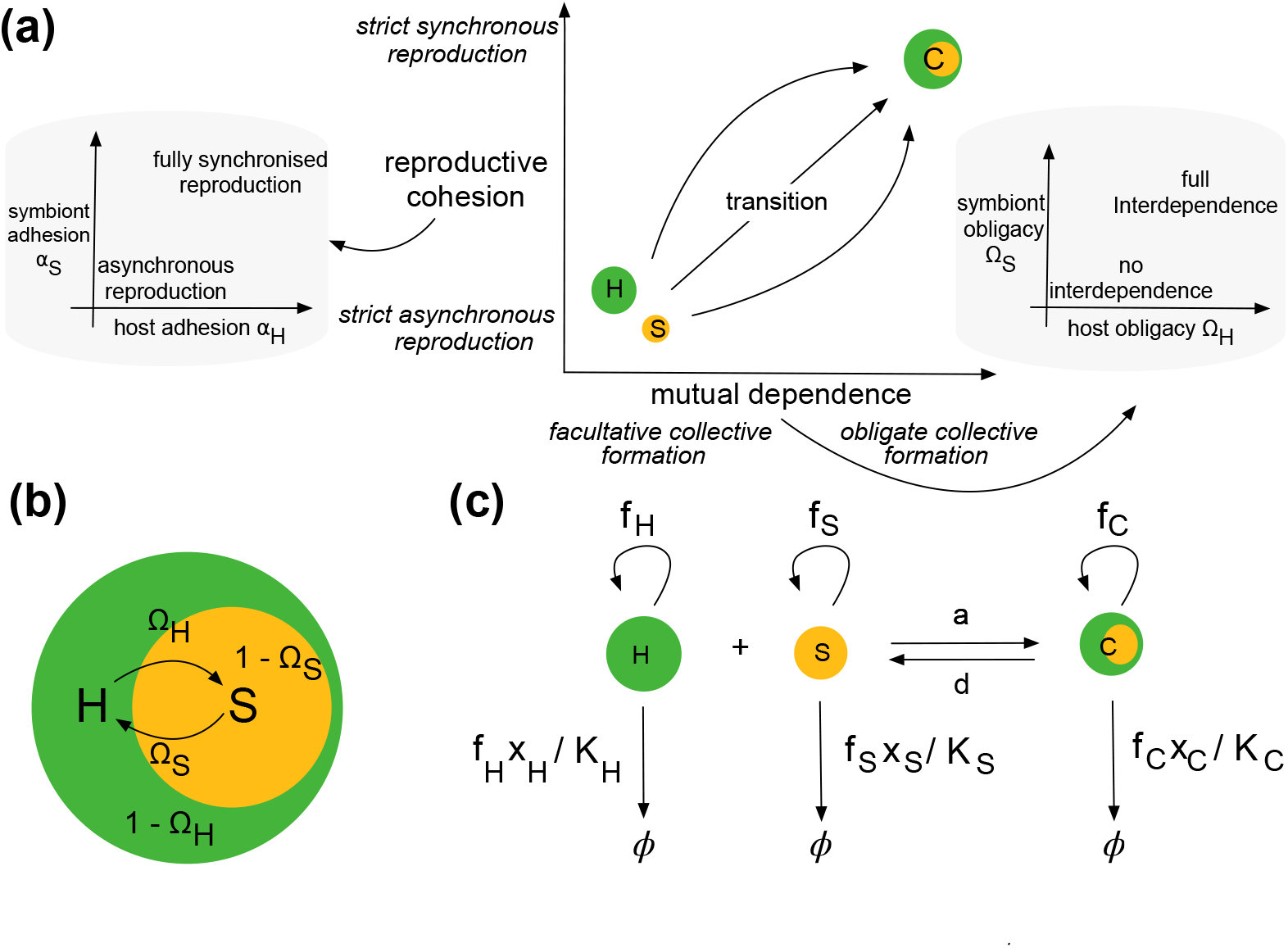
**(a)** We conceptualise the trajectory of an evolutionary transition in the plane of two quantities - the reproductive cohesion of the lower-level individuals and the degree of mutual dependence between them. The adhesions *α*_*H*_ and *α*_*S*_ of the host and symbiont respectively control their investment in reproductive cohesion, i.e., the degree of synchronised reproduction. The obligacy Ω_*i*_ describes the investment of type *i* in growth as a collective. **(b)** On a microscopic scale, it is helpful to picture that there is – exclusively when they are both part of the collective – resource exchange between the host and symbiont. In this setting, the traits Ω_*i*_ control how much resource sharing the host/symbiont individuals are prone to. The trait *α*_*i*_ is more phenomenological and is most concretely connected to the magnitude of synchronised vs. asynchronised reproduction of species *i*. **(c)** The flows of the population dynamical model. All populations have a logistic growth rate corresponding e.g. to an intrinsic birth rate *f*_*i*_ and a density-dependent death rate *f*_*i*_*x*_*i*_*/K*_*i*_. The host and symbiont associate and dissociate with rates *a* and *d* respectively.

We formalise the ecological, i.e. population dynamics over short, mutation-free timescales as follows: Let *f*_*H*_, *f*_*S*_, *f*_*C*_ be the growth rates of the host, symbiont, and collective populations, respectively. Further, suppose the host and symbiont associate with and dissociate from each other at rates *a* and *d*, flowing into and out of the collective population. See Fig. 1(c) for a graphical representation of all possible events. The dynamics are given by:

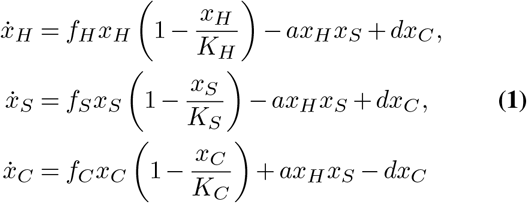

where *K*_*i*_ is the carrying capacity of population *i*. These dynamics are our default model, which we study extensively. To achieve some analytical results, we will also study this model with an infinite collective carrying capacity (*K*_*C*_ *→ ∞*). We also explore numerically different extensions, which we specify in the respective results sections.

The parameters of the population dynamical model above depend on the traits underlying dependence and cohesion. Consider first mutual dependence: we introduce the traits Ω_*H*_ and Ω_*S*_, referred to hereafter as the “obligacy” of the host and symbiont, respectively (Fig. 1a). These are dimensionless numbers in [0, 1] and denote the degree of dependence of the host and symbiont on the formation of the collectives. These traits formalise a tradeoff between individual (*f*_*H*_, *f*_*S*_) and collective reproduction rate *f*_*C*_. To make explicit the colocalised nature of an endosymbiotic interaction, we assume that the benefits of endosymbiosis are only present when the organisms are part of the collective, implying, e.g. that the growth rates of the host population in isolation do not depend on the symbiont’s investment (and vice versa).

With regards to reproductive cohesion, we introduce the traits *α*_*H*_ and *α*_*S*_, hereafter referred to as the “adhesion” of the host and symbiont, respectively. These are also dimensionless in [0, 1], and a higher adhesion denotes a higher association rate *a*, lower dissociation *d*, and a higher propensity of synchronised birth *f*_*C*_. These traits induce a tradeoff between processes favouring the formation of the collective and those favouring the individuals. The mathematical translation of these statements is in terms of the partial derivatives of ***f***_*i*_, *a, d* along the traits and is stated precisely in section §S.1.1 of the Supplementary Information (hereafter “SI”). The mapping of traits to population dynamical parameters that we will use for most of this work is

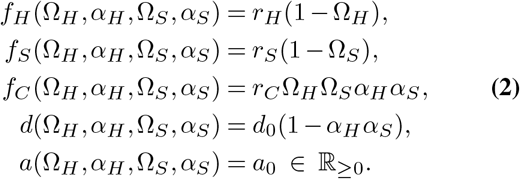

The parameters in our model are summarised in Table 1. In the Results section, we present analyses of a set of interconnected versions of the model set up thus far. We first study the evolution of dependence only – letting obligacies Ω_*i*_ evolve while the adhesions *α*_*i*_ stay constant – and then vice versa, i.e. cohesion only. We then introduce a simplified version of the population dynamical Eqs. 1 where the collective can grow exponentially; here we gain some analytical insights and study the joint evolution of obligacies and adhesions.

**Table 1.** A list of and associated information regarding all parameters named in our model.

| Parameters | Description | Possible values | Comments |
| --- | --- | --- | --- |
| $\Omega_H, \Omega_S$ | Host and symbiont obligacies | $[0,1]$ | initially set to $(0,0)$ |
| $\alpha_H, \alpha_S$ | Host and symbiont adhesions | $[0,1]$ | initially set to $(0,0)$ |
| $x_H, x_S, x_C$ | Population densities of host, symbiont, and collectives | $[0, \infty)$ | equilibrium densities depend on trait values |
| $f_H, f_S, f_C$ | Growth rates of H, S, and C populations | $[0, \infty)$ | usually set such that $f_S > f_C > f_H$ |
| $K_H, K_S, K_C$ | Carrying capacities of H, S, and C populations | $[0, \infty)$ | usually set such that $K_S > K_C > K_H$ |
| $a, d$ | Association and dissociation rates | $[0, \infty)$ | initial $d > \text{initial } a$ |
| $d_H, d_S$ | Within-collective mortality rates of host and symbionts | $[0, \infty)$ | assumed to not depend on any traits |
| $b_H, b_S$ | Within-collective birth rates of host and symbionts | $[0, \infty)$ | assumed to not depend on any traits |
| $r_H, r_S, r_C$ | Maximum values of $f_H, f_S, f_C$ | $[0, \infty)$ | Only relevant when $\Omega_i$ and/or $\alpha_i$ are evolving |
| $d_0$ | Maximum value of $d$ | $[0, \infty)$ | Only relevant when $\alpha_i$ are evolving |

We assume a growth tradeoff: that any investment in independent reproduction from either host or symbiont comes at a cost to their contribution to the collective growth rate (and vice versa). Notice that in Eqs. S3, host independent growth rate *f*_*H*_ is maximum at Ω_*H*_ = 0, decreases with increasing Ω_*H*_, and is zero at Ω_*H*_ = 1 (and identical for the symbiont). This tradeoff is inspired by the consideration that the selective pressures while in a collective and those while free-living are sufficiently contrasting that mutations have antagonistic effects in these two niches. For example, endosymbiont adaptation can be influenced by external pH levels or ambient amino acid availability, which are plausibly vastly different inside and outside the host. Similar statements about nutritional availability or pathogen defense can also be made for the pressures on host adaptation. Such tradeoffs have been observed in the squid-*Vibrio* symbiosis (41, 42), and in some experiments with *Bradyrhizobium* symbionts of plants (43) (see (44) for a more extensive discussion of growth trade-offs). We are in this study interested primarily in mutualistic endosymbiosis, where increased investment from either partner increases the collective growth rate. The tradeoff above is a case where investment into collective growth is a priori difficult to emerge because of the linked cost to free-living growth. It is possible that only the symbiont experiences such a tradeoff or that there are mutations that only advantage the collective or disadvantage the free-living host/symbiont; we do not consider these in our model.

To differentiate between the Ω and *α* trait pairs, it is useful to focus on an example such as a fig-wasp mutualism (reviewed in (45)). This association is an enmeshing of two life cycles – the fig tree depends on the wasp, the pollinator, and the wasp is dependent on the fig tree (the fruit, to be specific) to complete a part of its development. Therefore, the two species are highly dependent on each other, but they do not physically reproduce as a unit. Hence, here the obligacies Ω_*i*_ are high, but the adhesions *α*_*i*_ are not.

### Evolutionary invasion analysis

To compute an arbitrary mutant’s invasion fitness, we suppose a resident population is at ecological equilibrium, i.e., a stable steady state of the population dynamics (in our case Eq. (1)). The abundances at this steady state are computed numerically, unless mentioned otherwise. It is assumed that a mutant then arises, with trait value drawn from a symmetric probability distribution centred at the resident trait. The fate of this mutant is decided via its invasion fitness, i.e. the growth rate of a small number of mutants in the resident population (37). The mutant can invade if its invasion fitness is positive; if it is negative, the mutant goes to extinction (38, 46). This quantity is traditionally defined as the largest eigenvalue of the Jacobian of Eq. (1) when augmented for the presence of a mutant type. However, in our case, this quantity is not amenable to mathematical analysis, so we use the next-generation theorem, which gives an alternate characterisation of the same number (47, 48). We then use the canonical equation of adaptive dynamics (38) to study the macroscopic behaviour of long-term evolutionary trajectories (SI §S.4).

## Results and Discussion

In the first subsection, we investigate the case where obligacies evolve independently of the adhesions and vice versa. In the second subsection, we analyse an extension of the model that includes within-collective birth and death rates. Then, in the third subsection, we study the relative importance of the collective population’s growth rate and carrying capacity in deciding the course of evolution. Finally, in the last subsection, we study obligacy and adhesion coevolution in a model more amenable to analytics, which we derive by assuming an infinitely large collective carrying capacity in Eqs. 1.

It is challenging to analytically solve the system of Eqs. Eq. (1) to determine the fixed points and their stability. However, we determined computationally that for a wide range of parameter values, the dynamics converges to a stable fixed point over the evolutionary change in our traits of interest (see SI §S.2.1). Moreover, the values of the equilibrium population sizes increase or decrease gradually with the trait values, suggesting that there is a single internal fixed point. We will assume throughout that *K*_*S*_ *> K*_*H*_ and *r*_*S*_ *> r*_*H*_, i.e. the symbiont population has a higher carrying capacity and reaches it faster than the host. Consequently – on average over a given duration of time – more symbiont mutants arise than host mutants, and the symbiont trait hence has a higher rate of evolution (assuming, as we do, that the rate of mutation and the variance in the mutant trait distribution are the same for host and symbiont traits; see Eq. 4.12 in (38)).

### Independent evolution of obligacies and adhesions

We begin by understanding the evolution of the obligacies when adhesions are kept constant, and vice versa. The mapping of obligacies to population dynamical parameters we use in Fig. 2, panels (a,b,c) is:

**Fig. 2.**
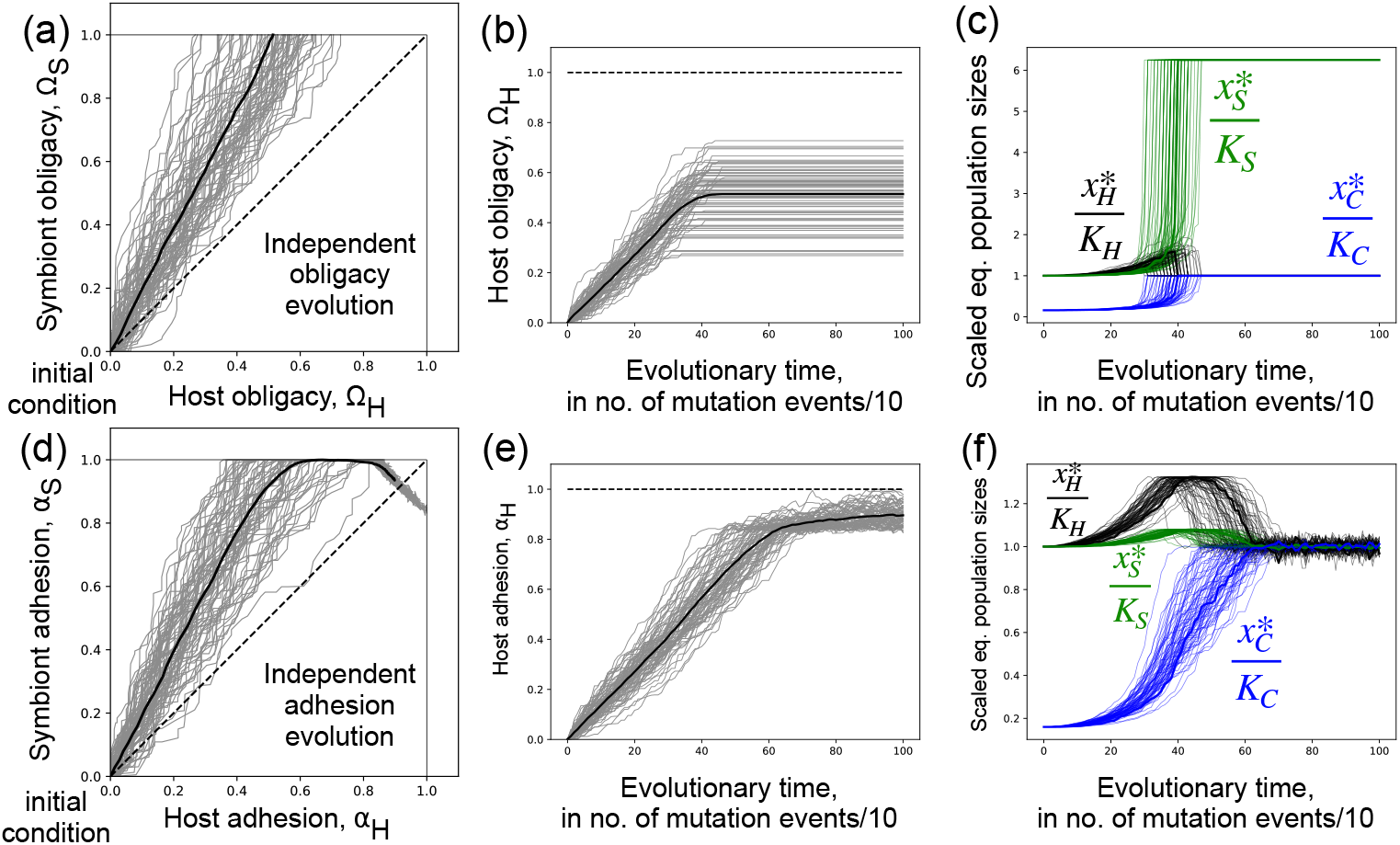
Evolutionary dynamics of host and symbiont obligacies. All panels contain results of 75 independent stochastic simulations. **(a)** Symbiont obligacy evolves to 1, and then host obligacy is no longer under selection (see main text for an explanation). **(b)** Symbiont adhesion evolves to 1, and then host adhesion increases until a certain value. The traits together then vary along a neutral ridge around (1,1) (see main text for an explanation). **(c**,**d)** These panels show the evolutionary trajectory of the host trait, since it is here that the nontrivial outcomes take place. Ω_*H*_ plateaus at different values based on when the value of Ω_*S*_ reaches one. *α*_*H*_ increases up to a high value and stays close to it. **Parameter values**. Common for all: *K*_*H*_ = 100, *K*_*S*_ = 200, *K*_*C*_ = 250, *a*_0_ = 0.1; for **(a)**: *r*_*H*_ = 8, *r*_*S*_ = 20, *r*_*C*_ = 10, *d* = 50.0; for **(b)** and **(c)**: *f*_*H*_ = 8, *f*_*S*_ = 20, *r*_*C*_ = 10, *d* = 50.

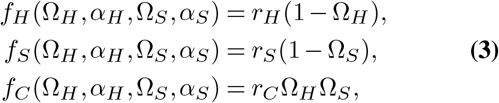

where we assume that the adhesions *α*_*i*_ are not under selection. The constants *r*_*H*_, *r*_*S*_, *r*_*C*_ set the scale of the parameters *f*_*i*_; concretely, they can be understood as the intrinsic growth rates of the populations at either Ω_*i*_ = 0 (for *f*_*H*_ and *f*_*S*_), or when both obligacies are equal to 1 (for *f*_*C*_). In panels (d,e,f) of Fig. 2, we assume obligacies to be constant and model the ecological consequences of a different adhesion via:

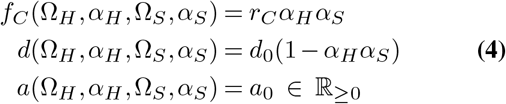

The constants *r*_*C*_, *d*_0_, *a*_0_ are again the maximum values of the respective functions, and set the scale of variation.

Results are presented in Fig. 2. First consider the evolutionary trajectory of the obligacies (Ω_*H*_ (*t*), Ω_*S*_(*t*)): the symbiont obligacy Ω_*S*_ reaches 1 first (denote this time by 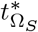), and after this the selective pressure on Ω_*H*_ disappears, leading to stagnation in its value (Fig. 2a,b). This lack of selection after Ω_*S*_ = 1 occurs because when *f*_*S*_ = 0 (no independent symbiont reproduction), the host and collective populations equilibrate to their carrying capacities irrespective of Ω_*H*_ ; the symbiont population is sustained only through dissociation (see Eq. (1)). However, the symbiont has a higher equilibrium population size when Ω_*S*_ = 1 than when Ω_*S*_ *<* 1 due to the “backflow” from the dissociation of the collectives (see SI §S.2.1, or from Eq. (1)). The converse is valid for the host – its population size is not at the maximum possible over the trait space, while Ω_*H*_ *<* 1. This is a counterintuitive result: the symbiont foregoes its ability to reproduce independently and has a higher equilibrium population size than its carrying capacity; the host now has a population size exactly equal to its carrying capacity (when it could be higher, i.e. at Ω_*H*_ = 1), but retains its ability to reproduce independently (Fig. 2c). Indeed, it is better (in terms of abundance) for both the host and symbiont to give up independent reproduction and be sustained only through dissociation, but the faster-evolving population can do this first. The above argument can also be made analytically via the computation of the basic reproductive number of a mutant 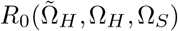, which denotes – roughly – the number of offspring left by a mutant host with obligacy 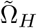 that arises in a resident population of hosts with obligacy Ω_*H*_ and symbionts with obligacy Ω_*S*_ (derived using the next-generation theorem, see SI §S.4.1). This quantity is given by

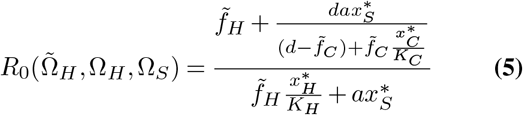

where a star denotes that the value at equilibrium must be used. The *R*_0_ is not maximally informative in our case since we cannot analytically solve for the equilibrium population abundances 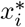. Nevertheless, it can be shown (see §S.4.1) that 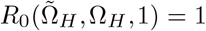 for any combination 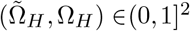 i.e. there is no fitness difference between any Ω_*H*_ mutants that arise in the background of Ω_*S*_ = 1. A consequence of this result that cannot be captured by adaptive dynamics is that this model predicts the accumulation of neutral genetic variation in the host population at loci coding for obligacy.

Now consider the evolutionary trajectory of adhesions, presented in Fig. 2d,e. Symbiont adhesion *α*_*S*_ reaches 1 first while *α*_*H*_ is still smaller than 1, and then host adhesion increases until an evolutionarily stable strategy (ESS) is reached. Trajectories then drift along a curve near (1,1). This ridge connects the states (Ω_*H*_ *<* 1, Ω_*S*_ = 1) and (Ω_*H*_ = 1, Ω_*S*_ *<* 1), and it is possible that a small proportion of trajectories eventually reaches Ω_*H*_ = 1. The drift in Fig. 2d,e,f is what our model predicts of biological populations, however its source in this figure is different: accepting false positives when deciding the fate of mutants based on *R*_0_ *>* 1, which arise due to floating point errors that we cannot entirely remove. All neutral mutations would have *R*_0_ = 1, whereas we accept values only larger than a value that is infinitesimally larger than one to exclude such errors (see associated scripts for exact details).

Nevertheless, there are two facts to explain: that the traits increase and that there is a host-symbiont asymmetry in outcomes. Our observations can be explained by the variation in equilibrium population sizes as traits change (see SI §S.2.1 for heatmaps). Notably, (i) they increase along the adhesions *α*_*i*_ (and hence the traits themselves also increase); and (ii) they do so non-monotonically, with a local maximum in 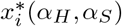 occurring over a ridge in the interior of the *α*_*H*_ − *α*_*S*_ space that surrounds the top right corner (1,1) (see Fig. §S.2.1). The first consequence is that any trajectory (*α*_*H*_, *α*_*S*_) must pass over this ridge to reach (1, 1). The symbiont adhesion can reach 1 before encountering this bump because of its faster rate of change, and the trajectory has to overcome it then while *α*_*H*_ is still less than 1. This is the source of the asymmetry: If both species have the same *r* and *K*, then the landscape of 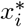 is the same, but now the trajectory (*α*_*H*_, *α*_*S*_) hits the ridge when both adhesions are approximately equal and less than 1 (see SI §S.3.1). We suggest that the inflection point of this local maximum corresponds to the ESS, since decreasing population size (even locally) is disadvantageous, and such a mutant could not invade.

More proximately, an increased adhesion leads to modifications in two forces acting on host/symbiont population size via the term *dx*_*C*_: (1) a decrease because fewer dissociations occur, but also (2) an increase because an increased adhesion increases *f*_*C*_, and hence the collective population size. Since we begin with the intuitive scenario of high dissociation rate and zero synchronised growth, an initial increase in adhesion is beneficial, but this benefit is only up to a point, after which the decrease of *d* overpowers the increase due to *f*_*C*_. This switch is the explanation for the presence of the intermediate ESS and the ridges in population size curves. This demonstrates that it is essential to consider not only intuitive parameters like *f*_*H*_ or *f*_*S*_ but also population size dynamics, which can strongly influence invasion fitness in this context.

### The effect of within-collective mortality and reproduction

In our model, we have assumed so far that dissociation is perfect: all events producing free-living individuals from a collective give rise exactly to one host and one symbiont. However, it is likely that there is differential mortality for hosts and symbionts while they are in a collective or during dissociation. This would give rise to free-living hosts when the symbiont dies (at some rate *d*_*S*_), and free-living symbionts with an analogous rate *d*_*H*_. Further, hosts and symbionts, while in a collective, can also give birth to free-living hosts or symbionts. Suppose this takes place at rates *b*_*H*_ and *b*_*S*_, respectively. The population dynamics is then described by

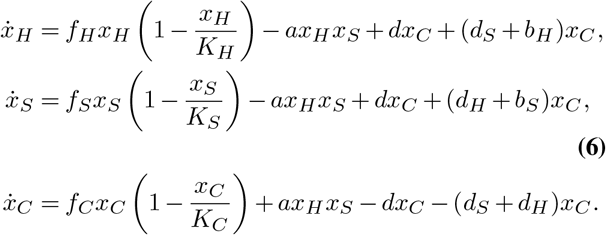

The previous section then corresponds to, e.g. setting the rates *b*_*i*_ = 0 and *d*_*i*_ = 0 *i ∈* {*H, S*}. The correspondence is slightly more general; one obtains Eqs. 1 whenever *b*_*i*_ = *d*_*j*_ for all combinations of *i, j∈* {*H, S*} ; with the caveat that the corresponding value of the dissociation rate would be higher, i.e. *d* + *b*_*i*_ + *d*_*j*_.

We consider two scenarios: (i) differential mortality while in collective, but no within-collective reproduction (*d*_*H*_ = *d*_*S*_ ≠ 0, *b*_*i*_ = 0 *i∈* {*H, S*} ; Fig. 3 right), and (ii) within-collective H (or S) can give birth to free-living H (or S) but within-collective mortality is absent (*b*_*H*_ = *b*_*S*_ ≠ 0, *d*_*i*_ = 0 *i ∈* {*H, S*}; Fig. 3 left). For the sake of brevity, we do not consider the cases where *b*_*H*_ ≠ *b*_*S*_ or *d*_*H*_ ≠ *d*_*S*_, or where the within-collective birth and death rates are both greater than zero. However, the latter is not modelling a completely new phenomenon since *d*_*i*_ and *b*_*j*_ appear together as an aggregate coefficient of the collective abundance *x*_*C*_ in Eq. (S14). First, the adhesion trajectories (Fig. 3 right): there are no qualitative changes compared to the case when *d*_*i*_ = *b*_*i*_ = 0, the prediction that the adhesions increase and then hit a ridge of evolutionarily stable strategies is robust to the addition of the above within-collective rates. At higher shared values of *d*_*i*_, the adhesions do not increase from initial values (see SI §S.3.4).

**Fig. 3.**
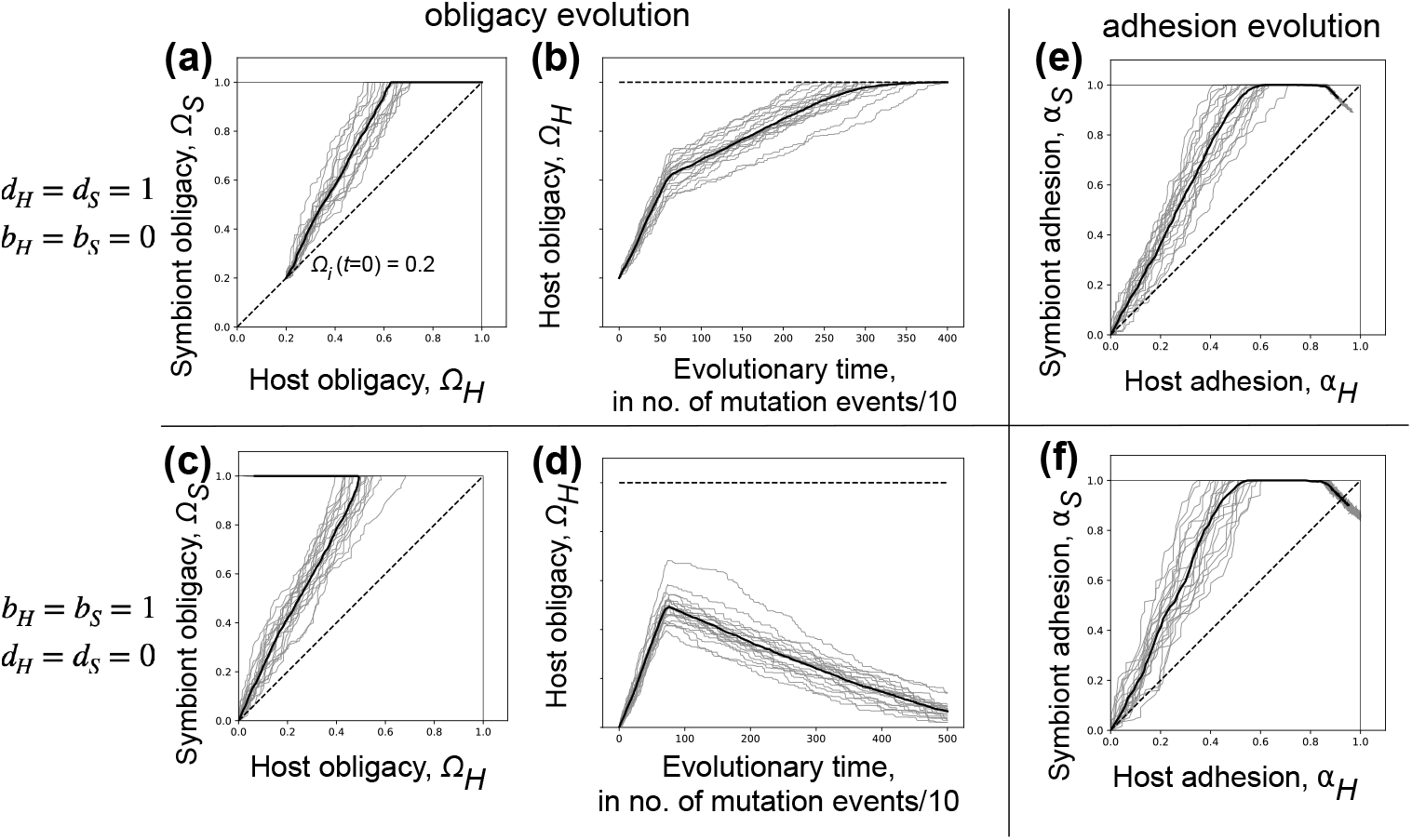
Evolutionary dynamics of obligacies and adhesions when *b*_*i*_ and *d*_*i*_ are nonzero. All panels contain results of 20 independent stochastic simulations. **(a**,**b)** When *d*_*H*_ = *d*_*S*_ *>* 0, the initial obligacies must be above a threshold for them to further increase. Symbiont obligacy increases faster, and when Ω_*S*_ hits 1, host obligacy increases further upto 1. **(c**,**d)** When *b*_*H*_ = *b*_*S*_ *>* 0, dependence directly increases over time without a threshold minimum value that is necessary; when Ω_*S*_ hits 1, host obligacy now decreases to zero. **(e**,**f)** Qualitatively identical to adhesion evolution presented in Figure 2. **Parameter values**. Common for all: *K*_*H*_ = 100, *K*_*S*_ = 200, *K*_*C*_ = 250, *a*_0_ = 0.1. Panels **(a**,**b**,**e)**: *d*_*H*_ = *d*_*S*_ = 1, *b*_*H*_ = *b*_*S*_ = 0. Panels **(c**,**d**,**f)**: *d*_*H*_ = *d*_*S*_ = 0, *b*_*H*_ = *b*_*S*_ = 1. Panels **(a**,**b**,**c**,**d)**: *r*_*H*_ = 8, *r*_*S*_ = 20, *r*_*C*_ = 10, *d* = 50.0. Panels **(e**,**f)**: *f*_*H*_ = 8, *f*_*S*_ = 20, *r*_*C*_ = 10, *d*_0_ = 50.

The evolution of obligacies, however, displays departures both when *d*_*i*_ *>* 0 or *b*_*i*_ *>* 0. When hosts or symbionts can die while in a collective but cannot give birth, the obligacies do not initially increase from (0,0) at all. However, if the obligacies are at a high enough initial value to begin with, the obligacies then increase as before, with the symbiont obligacy Ω_*S*_ reaching 1 first. This threshold initial value increases with the shared value of *d*_*H*_ and *d*_*S*_, but above this value the dynamics proceed similarly to the earlier model (see SI §S.3.4), with one important difference: once Ω_*S*_ = 1, the selective pressure on Ω_*H*_ does not disappear, instead now the host obligacy also increases to 1, leading to full mutual dependence. At the state Ω_*H*_ = Ω_*S*_ = 1, the equilibrium population abundances remain at steady, non-zero values in our case of *d*_*S*_ = *d*_*H*_.

When hosts and symbionts can give birth while in a collective but cannot die, the outcome is again different: the obligacies directly increase from (0,0) without needing a threshold dependence, but once symbiont obligacy reaches 1, the host is under selection to now decrease its obligacy. This leads to a non-monotonic route to a one-sided mutualism, where the symbiont is completely dependent and the host not at all. In the ultimate state of Ω_*H*_ = 0, Ω_*S*_ = 1, there is no fixed point possible for Eqs. S14; the symbiont goes extinct, while the host and collective populations blow up due to the birth rates. At higher shared values of *b*_*i*_, the obligacies still increase, but host-symbiont symmetry in obligacies is reduced (see SI §S.3.4).

To explain the results above, we again appeal to the population sizes at equilibrium. It can be shown that, if *d*_*S*_ + *b*_*H*_ = *d*_*H*_ + *b*_*S*_ (which is true in Fig. 3 by the parameters we have set), once Ω_*S*_ = 1 (implying *f*_*S*_ = 0), the equilibrium abundances calculated from Eqs. S14 are given by the equations

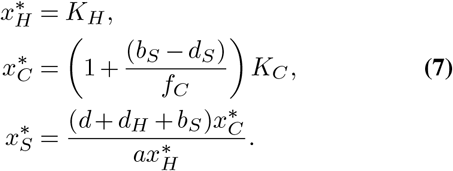

Dependence on Ω_*i*_ here enters through the *f*_*C*_ in 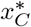: when *b*_*S*_ *> d*_*S*_, an increased host obligacy decreases collective abundance at equilibrium 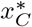, and when *b*_*S*_ *< d*_*S*_, an increased host obligacy *increases* 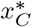. More generally, the form of 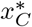 reflects that the forces driving the collective population away from its carrying capacity are *b*_*S*_-produced free-living symbionts which form collectives via host association, and collectives leaving the compartment at rate *d*_*S*_ to form hosts. This suggests that even if the rates *b*_*i*_ and *d*_*i*_ are unequal, a case we have not tested here, the difference *b*_*S*_ − *d*_*S*_ plays a central, if not solitary, role in deciding the outcome.

The inclusion of births and deaths of hosts and symbionts while in collective therefore affects two properties of the trajectory: initial increase of the obligacies, and what happens when Ω_*S*_ = 1 is reached (see Table 2 for a summary of these effects). Including trait-dependence onto these within-collective rates would further complicate the picture since their values would become dynamic, and combinations of the above arguments might become necessary. We do not consider this possibility here, but some possible consequences are discussed in the Conclusions section. Having clarified the effects of the parameters *b*_*i*_ and *d*_*i*_, we shall set them again to zero for the remaining sections to emphasize other predictions of this model.

**Table 2.** A summary of evolutionary outcomes for Ω_*i*_ and *α*_*i*_ depending on the rates of within-collective mortality and reproduction.

| Scenario | Parameters | Consequences on $\Omega_i(t)$ | Consequences on $\alpha_i(t)$ |
| --- | --- | --- | --- |
| Perfect collective dissociation, no within-collective births or deaths | $d_H = 0, d_S = 0$<br>$b_H = b_S = 0$ | both increase, host obligacy stagnates once symbiont completely obligate | increase, then drift along a ridge |
| Within-collective mortality, but no births | $d_H = d_S > 0, b_H = 0, b_S = 0$ | both increase if initial obligacy high enough, ultimately both host and symbiont both become obligately dependent | increase, then drift along a ridge |
| Within-collective hosts can give birth to new hosts (similarly symbionts), but no within-collective deaths | $b_H = b_S > 0, d_H = 0, d_S = 0$ | both increase without a threshold, ultimately symbiont becomes fully obligately whereas host completely independent | increase, then drift along a ridge |
| Within-collective birth and death rates are all equal | $b_H = b_S = d_H = d_S$ | equivalent to setting $b_i = d_i = 0$, and increasing $d$, so there is no formal difference to the first row | |

### Collective growth rate and collective carrying capacity have different effects on evolutionary outcomes

We have shown that dependence and cohesion can, in fact, increase under our assumptions, and that associating with a partner to form a collective is thus beneficial. However, this benefit is in terms of two independent (at least in our model) parameters - the collective population’s growth rate *r*_*C*_ and its carrying capacity *K*_*C*_. In this section, we are interested in which of these parameters is more important in determining the evolutionary outcome. We computed evolutionary trajectories for high and low values of the parameters *r*_*C*_ and *K*_*C*_ each – *r*_*C*_ *∈* {1, 40} and *K*_*C*_ *∈* {10, 500}. At the low value of *K*_*C*_, both obligacies and adhesions remained at their initial condition (0,0); at the high value of *K*_*C*_, they evolved away from (0,0). The outcome was not influenced by the value of *r*_*C*_. This shows that it is more important that the collective’s equilibrium population size is high, no matter how much the mutant and resident differ in how fast they reach this size. We propose a cutoff for the threshold value of *K*_*C*_ in the following paragraphs.

Nevertheless, once *K*_*C*_ is high enough, its precise value can influence the ultimate values of adhesion, but not obligacy. Fig. 4 presents the results of evaluating the evolutionary trajectories of obligacies and adhesions at two values of the parameters *K*_*C*_, both of which are high enough to allow traits to increase from (0,0) but quantitatively different.

**Fig. 4.**
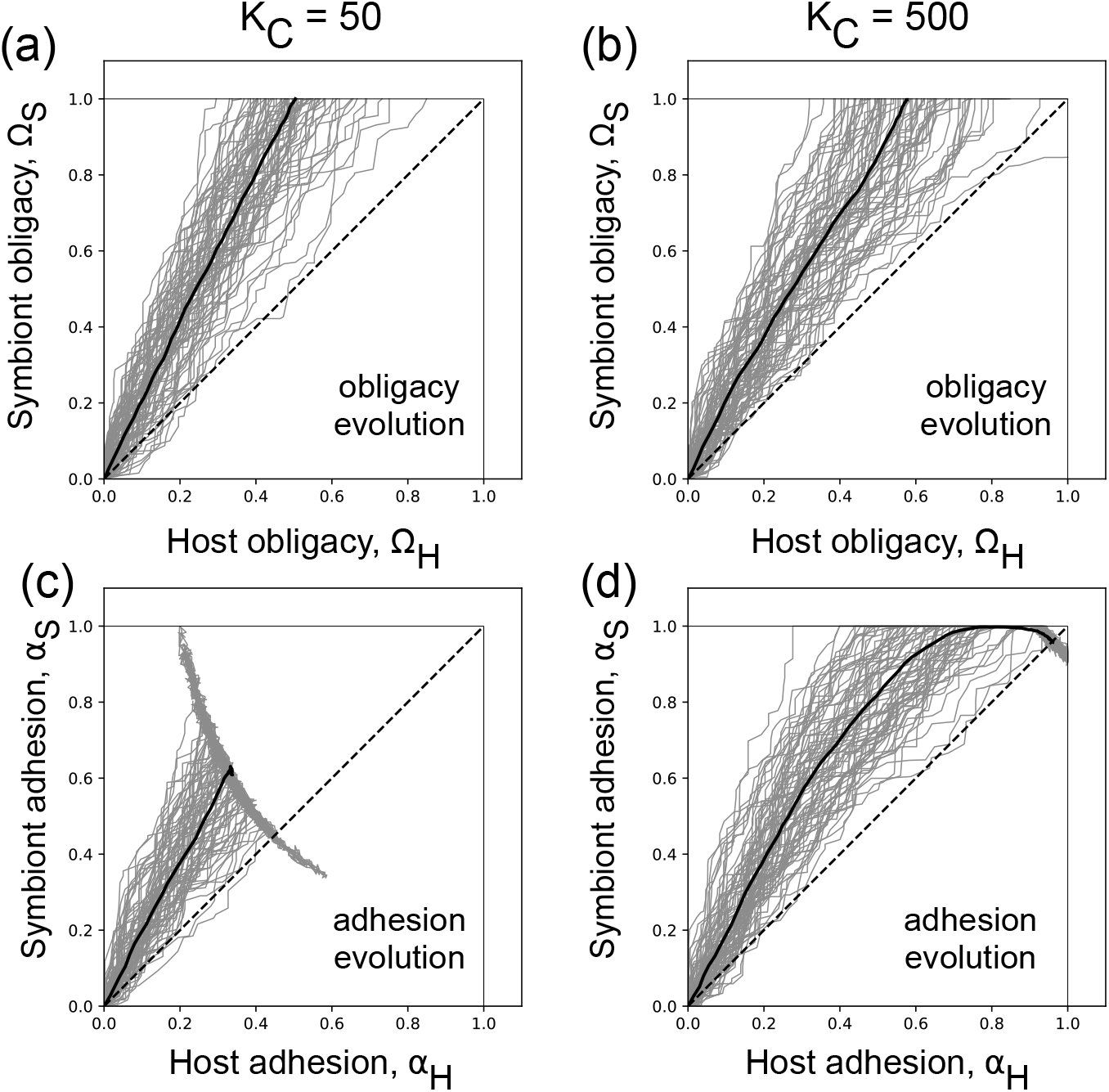
Collective carrying capacity qualitatively and quantitatively affects ultimate trait values. Each panel contains the results of 75 independent simulation runs (in grey), with the average trajectory in solid black. Both values of *K*_*C*_ = 50, 500 in this figure are high enough to allow traits to increase from (0,0).*K*_*C*_ = 10 was not. When *K*_*C*_ is high enough, obligacies increase such that symbiont obligacy evolves to 1 and host obligacy subsequently is no longer under selection. Ultimate values of adhesion are affected by the precise values of *K*_*C*_. The question of how high *K*_*C*_ must be is addressed in the main text. This outcome did not change between simulations run under low and high values of *r*_*C*_ = 1 and *r*_*C*_ = 40 respectively. **Parameter values**. *a*_0_ = 0.1, *d* = 50, *r*_*H*_ = 8, *r*_*S*_ = 20, *r*_*C*_ = 40, *K*_*H*_ = 100, *K*_*S*_ = 200; *r*_*C*_ and *K*_*C*_ vary as indicated in the figure.

Obligacy evolution is not affected by the value of *K*_*C*_. This is in line with our intuition developed in the previous sections – the defining event in an increasing obligacy trajectory is the location of its arrival at the unit square’s boundary, and this cannot be affected by *K*_*C*_, a parameter that does not bias trajectories preferentially towards hosts or symbionts. The bias Ω_*S*_ *>* Ω_*H*_ does indeed persist, induced by the difference in host and symbiont population growth parameters – something we did not manipulate for these tests.

Adhesion trajectories, however, are affected by the precise value of *K*_*C*_. This is because the important event here is the trajectory’s arrival at the neutral ridge of ESSs described in the previous section. The presence of this ridge is explained by the non-monotonically varying benefit that dissociation of collectives provides to independent host and symbiont populations. Therefore, it is intuitive that the ridge’s location is affected by *K*_*C*_, a parameter that strongly sets the equilibrium abundance of collectives. At lower *K*_*C*_, the relative number of collectives is lower and hence dissociation becomes non-beneficial at relatively lower values of the adhesions and hence relatively higher values of the effective dissociation rate *d*_0_(1 − *α*_*H*_ *α*_*S*_).

Note that the ridge when *K*_*C*_ = 50 also prevents the adhesions from reaching maximum values even for the symbiont (despite the bias *α*_*S*_ *> α*_*H*_ persisting for identical reasons as above). When the carrying capacity *K*_*C*_ is very high (see SI §S.3.3), this ridge is extremely close to (1,1) and the adhesions effectively reach their maximum values. The slight movements around 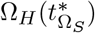 and the drift of adhesions along the ridge are artefacts of floating-point errors as before and can be ignored; we refrain from finding an involved artificial way to remove these since their origin is provably clear (see previous sections).

The *r* − *K* difference in importance is also reflected in the variation of the equilibrium population sizes as traits evolve (SI §S.2.1). The pattern of variation supports the claims we have made so far – it is not qualitatively affected by the value of *r*_*C*_, but is so affected by changes in *K*_*C*_. The ridge in population sizes can also be shown to be closer to (1,1) for higher *K*_*C*_. This agrees with the intuition that the scale of the equilibrium population size 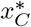 is set more strongly by *K*_*C*_, with a smaller spread around it that is determined by other parameters. The maximum population size attained by the collective is its carrying capacity, and this takes place when at least one of (Ω_*H*_, Ω_*S*_) = 1, and when both *α*_*H*_ = *α*_*S*_ = 1. Our analysis suggests that the pattern of variation switches depending on which quantity is bigger - *dK*_*C*_ or *aK*_*H*_ *K*_*S*_, i.e. outflow or inflow into the population *i* when Ω_*i*_ = 1. In particular, when Ω_*H*_ = Ω_*S*_ = 0, population sizes equilibriate to 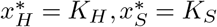; when Ω_*i*_ = 1 for *i* = {*H, S*}, then 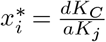 where *j* is the other species. This suggests a sufficient condition on how high *K*_*C*_ must be: if 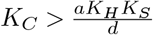, hosts and symbiont populations are incentivised to increase obligacy. We tested this by varying *a* and *d* to perturb the same threshold quantity, and the results support our claim (see SI section §S.3.2). These arguments are difficult to make fully explicit given intractable analytical results – a problem that we directly address in the next section.

### Does mutual dependence evolve before or after reproductive cohesion?

To understand the co-evolution of both obligacies and adhesions across the host and symbiont, we introduce a simpler version of Model Eq. (1). The motivation is two-fold: first, it allows for analytical tractability, enabling better understanding of our model; second, it restricts focus to the early evolutionary dynamics, since the dynamics at the boundaries is strongly dependent on the parameters *b*_*i*_ and *d*_*i*_. This simpler model assumes that the collective population exhibits unbounded exponential growth, i.e. that the carrying capacity is infinitely high. The “exponential model” will henceforth refer to the following system of ordinary differential equations:

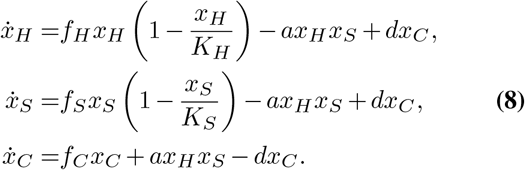

Analogously, Eq. system Eq. (1) will now be referred to as the “logistic model” when necessary. One must keep in mind that comparisons to the logistic model of the previous sections can be made only when the traits Ω_*i*_ and *α*_*i*_ (and hence *f*_*C*_, see the mapping Eq. (S3)) are of small value, since once *f*_*C*_ is high enough, the self-limitation term −*f*_*C*_*x*_*C*_*/K*_*C*_ of the logistic growth rate becomes relevant. The formation of the collective can here be formally shown to be comparable to a mutualism between the host and symbiont, see SI §S.1.2. Many statements can now be made regarding the existence, feasibility, and stability of population dynamical equilibria. The precise statements are relegated to the SI, in §S.2.2 and §S.4.2; however, the important features are that there is only one stable internal fixed point, which is feasible when the dissociation rate *d* is high enough to counteract the exponential growth of the collective. This is also borne out in the evolutionary dynamics of Ω_*i*_ and *α*_*i*_ under the exponential model (see §S.4.2); the traits increase until a point, and much before either has reached its maximum value, the collective’s reproduction is so strong that no ecological equilibrium exists and the collective population blows up to infinity. Analogous to Eq. (5), an *R*_0_ value can also be calculated here, and it is very close in functional form to the *R*_0_ under the logistic model. In particular, it is given by

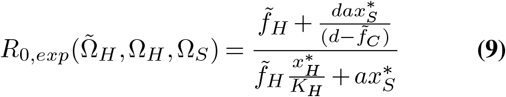

and is exactly the result of taking *K*_*C*_ *→ ∞* in the logistic *R*_0_ (see Eq. (5)). This shows that the evolutionary dynamics captured by the exponential model are, at early trait values, truly comparable to the logistic model Eq. (1). In fact, one can go further here, and sderive an exact invasion criterion – suppose a host mutant arises with obligacy 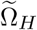 and adhesion 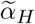. We show (§S.4.2) that this mutant will invade if and only if

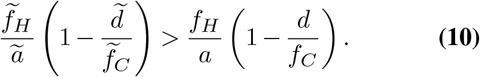

where quantities with a tilde (*∼*) are associated to the mutant. Since the host and symbiont are in our model identical in everything but the label we impose on them, an analogous criterion exists for the fate of a symbiont mutant. This criterion is “separable” into two quantities of the same functional form, each depending only on either the mutant or the resident. This implies that over the course of successive mutations, this underlying quantity is maximised. The existence of such an optimisation principle (49) also means that there can never be evolutionary branching in our traits at these early times, see §S.4.4. Finally, using the constraints we set up in the Model section and §S.1.1, it can also be shown that all four traits Ω_*H*_, Ω_*S*_, *α*_*H*_, *α*_*S*_ must monotonically increase over evolutionary time.

To study dependence-cohesion coevolution, we again consider the mapping in Eq. (S3); results of the simulations are shown in Fig. 5. We represent the degree of dependence between the host and symbiont by the product Ω_*H*_ Ω_*S*_ of their obligacies, and the degree of reproductive cohesion by *α*_*H*_ *α*_*S*_^1^. These results show two important facts. First, for both the host and symbiont, it is adaptive to evolve such that Ω_*i*_ *> α*_*i*_, i.e. being more obligate than cohesive, results from natural selection. Focusing on Ω_*H*_ Ω_*S*_ and *α*_*H*_ *α*_*S*_, one concludes that evolutionary trajectories are biased toward more mutual dependence than reproductive cohesion. This is a central result. It shows that, over time, we expect that a host-symbiont collective evolves such that the partners are closer to complete mutual dependence than to reproducing synchronously.

**Fig. 5.**
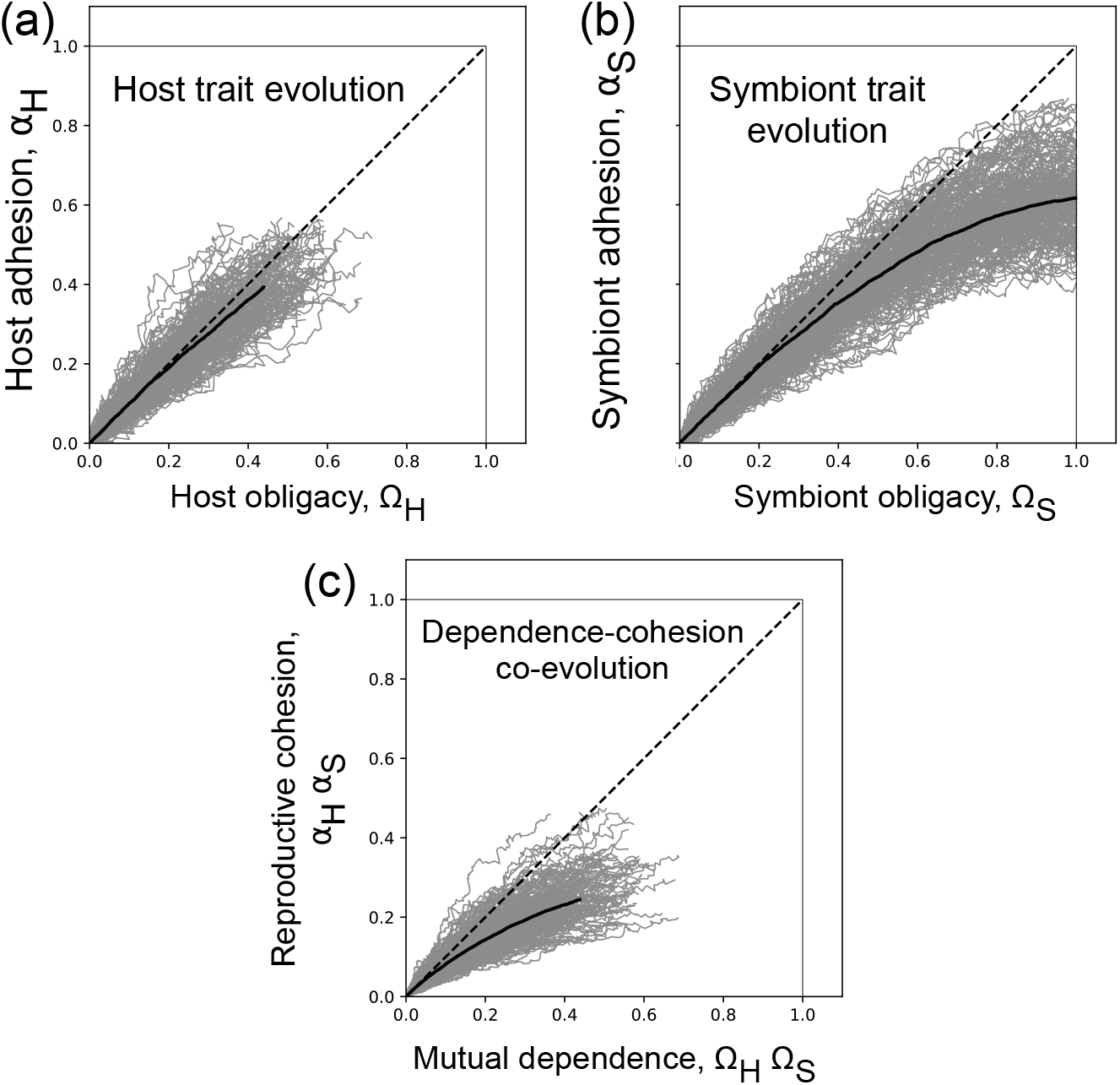
Mutual dependence evolves before collective reproduction. Here we consider the co-evolution of the four traits Ω_*H*_, *α*_*H*_ and Ω_*S*_, *α*_*S*_ under the exponential model (Eqs. S6). **(a, b)** Both host and symbiont traits evolve such that, on average, Ω_*i*_ *> α*_*i*_. Notice that symbiont evolution (panel **(b)**) shows that symbiont traits evolve to much higher values than host traits. **(c)** Given the individual evolutionary trajectories of the host and symbiont, one can collate information to obtain measures that describe the collective as a whole - the degree of mutual dependence is represented here by the product Ω_*H*_ Ω_*S*_ and the degree of reproductive cohesion by *α*_*H*_ *α*_*S*_. We observe that mutual dependence evolves faster than reproductive cohesion. The trajectories stop when they hit the feasibility bound; it is nontrivial to visualise its manifestation in this plane and is hence absent. **Parameter values**. *K*_*H*_ = 100, *K*_*S*_ = 200, *K*_*C*_ = 500, *a*_0_ = 0.1, *r*_*H*_ = 8, *r*_*S*_ = 20, *r*_*C*_ = 10, *d*_0_ = 50.0.

Fundamentally, this is because the functional effects of the pairs of traits are different. Recall that both trait pairs increase because of the better growth of the collective. However, there is a difference between them: independent growth of type *i* (*∈* {*H, S*}) is a function only of Ω_*i*_, whereas the effect of the *α*_*i*_ occurs only in “coordination” with both species since dissociation is a function of the product of *α*_*H*_ and *α*_*S*_ (see Eq. (S3)). Therefore, an increase in Ω_*i*_ has a higher functional effect (in increasing the invasion fitness) than the same increase in *α*_*i*_, since the latter’s effect is damped by the other adhesion as well (since *α*_*i*_ *∈* [0, 1]). See SI §S.4.5 for a more comprehensive explanation via other functional form choices.

This result is robust to the choice of parameter values, different generation times, additive (instead of multiplicative) effect of the individual traits on the collective, etc. (SI Fig. S.14). The bias persists, and mutual dependence evolves faster than reproductive cohesion. This strongly suggests that endosymbioses in nature are more likely to be mutually dependent than cohesive.

## Conclusions and conjectures

Endosymbiosis and the advances in complexity it made possible are astonishing. An endosymbiotic association led to eukaryotes and many other intricate associations between unrelated individuals. In this study, we endeavour to give a precise view of endosymbiosis as an egalitarian evolutionary transition in individuality and study the effect of some essential ecological factors on its origins.

We study two significant characteristics of an endosymbiotic collective undergoing an evolutionary transition - the reproductive cohesion of the host and symbiont (affected by host and symbiont adhesion, *α*_*H*_ and *α*_*S*_), and the level of mutual dependence between them (obligacy, Ω_*H*_ and Ω_*S*_). Our model shows that when obligacies evolve independently, one expects the symbiont obligacy to increase to its maximum value first, which can lead to the host obligacy being under selection to either increase, decrease, or stay constant under different ecological scenarios. This might explain the diversity in dependence outcomes that we see in nature, and we identify the two parameters in our model, *b*_*i*_and *d*_*i*_ (*i* = *H, S*), that control this outcome. The adhesions also increase (with symbiont adhesion reaching high values first), ultimately leading to the accumulation of neutral variation due to drift of the evolutionary trajectory along a line of evolutionarily stable strategies. The final value of the adhesions depends on the collective carrying capacity. Second, we show that host-symbiont asymmetries in evolutionary outcomes arise due to differences in their population growth rates and carrying capacities. Lastly, we show that at early evolutionary times, the density of permitted evolutionary trajectories in the dependence-cohesion plane is not uniform: when Ω_*i*_ and *α*_*i*_ co-evolve, both species evolve to be more obligate than adhesive, irrespective of the rate of these traits’ evolution.

Our work highlights the importance of considering differences between host and symbiont population growth parameters. That the symbiont trait (either obligacy or adhesion) increases faster than its host counterpart is due to the symbiont’s larger population size and hence faster rate of evolution: an instantiation of the Red King effect (50, 51) – such a bias must be expected when host and symbiont interests are aligned, and defection (lower Ω or *α*) leads to a lower payoff. In the original Red King effect, the interest-aligning mechanism is a mutualism – an ecological interaction. Our analysis shows that the collective’s formation (and shared fate) is another fundamental, interest-aligning mechanism in evolutionary transitions. Further, this faster rate of evolution has qualitatively novel effects in our model: in the case of obligacy, the symbiont evolves to full obligacy first, which leads to conditions where the host obligacy can be under selection to increase, decrease, or neutrally drift. Our model predicts full dependence for the interaction partner with higher growth rate and carrying capacity, and parameter-dependent scenarios for the partner with lower growth rate and carrying capacity. There are at least two more qualitatively different routes to asymmetric investments: first, the growth tradeoffs *f*_*H*_ (Ω_*H*_) and *f*_*S*_(Ω_*S*_) for host and symbiont investing in collective reproduction might be different, and might in nature take non-linear forms. The work of (52) suggests that this might have a qualitative impact: in their model, obligate exploitation can be observed over mutual dependence when the growth benefit of losing a costly function is accelerating in the amount of function lost. Further, we assumed that the within-collective reproduction and mortality rates are identical for host and symbiont. We assumed this for explanatory clarity, but it is almost certainly not true, and will give rise to new layers of asymmetry. Since these parameters are central to the evolutionary outcome, it is imperative in future work to measure them experimentally and characterise differences in their values. Developing this part of the theory might also help understand why current host-symbiont relationships are often biased towards the symbiont being much more obligate than the host. This is clearest in the evolution of tiny genomes in endosymbionts (31, 53, 54), and the ability to go through symbiont loss and replacements in some hosts (55). However, reductive genome evolution has also been shown to be driven by other mechanisms such as Muller’s ratchet (56) or environment-induced redundancy (57–59), in addition to Red King-type effects. These latter effects are predicated on endosymbionts occurring within the hosts and not the other way around. Simple factors such as growth parameters and the nested structure of endosymbioses are therefore clearly important, and further work is required to delineate their consequences from that of complicated strategies such as partner choice/sanction or zero-determinant strategies (60, 61).

Another main aim was to understand why an evolutionary transition does not always take place. In this context, our model suggests the importance of the within-collective birth and death rates. They control model behaviour at two important timepoints: whether the obligacies initially increase, and what happens to the host obligacy once the symbiont becomes completely obligate. If within-collective births are stronger than deaths, (i) both obligacies increase initially, and (ii) host obligacy decreases to zero once the symbiont has become obligate. If the opposite is true, i.e. within-collective deaths are stronger than births, (i) the obligacies increase only above a certain threshold initial value, and (ii) the host becomes completely obligate once the symbiont has done so. Our model thus demonstrates that the evolutionary dynamics of dependence can impose strong constraints on the emergence of obligate endosymbioses. If the within-collective rates depend on the evolving traits Ω_*i*_ and *α*_*i*_, it would allow for the situation where births are stronger than deaths initially (so both host and symbiont obligacies initially increase) and deaths are stronger than births later (and hence both become fully obligate). However, the rates may depend on different (combinations of) traits, and different dependence-structures of *b*_*i*_, *d*_*i*_ on Ω_*i*_, *α*_*i*_, *i ∈* {*H, S*} might have different consequences. The values of *b*_*i*_ and *d*_*i*_ also need not be equal in the way that we have considered. Understanding the consequences of these extensions constitutes, in our view, the next study necessary in this body of work. Lastly, full adhesions can be achieved only when the collective carrying capacity is very high; when it is lower, final adhesion values must be interpreted carefully since our model predicts drift over time and hence variability.

The prediction that dependence evolves faster than cohesion at early times can also be confronted with biological examples. Following the work of (25), we compiled a short, non-exhaustive list of well-studied endosymbioses (Table 3) where there is information on the level of vertical transmission (a proxy for reproductive cohesion) and the degree of mutual dependence. However, due to the qualitative nature of the data, the causal mechanics of the interactions are inconclusive. Some cases show direct connection to our theory: in the *Riftia*-*Endoriftia* endosymbiosis, there is high dependence but with horizontal transmission, others do not – the *Dictyostelium*-*Burkholderia* farming symbiosis is facultative from both sides and has a mixed mode of transmission (62). It is impossible to compare the degree of dependence and cohesion here. There is indirect evidence that mutual dependence is easier to evolve than reproductive cohesion, since it is widely observed empirically (63), understood well via e.g. the Black Queen effect (58), can evolve rapidly in an experiment (64), and in principle requires very few traits (52). On the other hand, few studies have precisely estimated quantities related to the level of vertical and horizontal transmission (65, 66). Further, it is not clear if there are genetic constraints that would lead to correlations in obligacy and adhesion – a strong enough correlation could shift in a new direction the bias towards codependence that we observe. On the theoretical side, an interesting extension of our model is to understand if the bias towards dependence that we see survives to later evolutionary times, when traits are closer to their final values. This study and its limitations thus highlight the need for a tighter connection between empirical work and theory and the requisite experiment where the complete evolutionary transition can be quantified. Nonetheless, taken as a null model, our study shows that an evolutionary transition in individuality is far from an inevitable outcome of the existence of a higher level of selection, suggesting that one might need to invoke different mechanisms to explain phenomena such as eukaryogenesis. While our model already allows for many of the scenarios that one would like to capture, many possible extensions exist that could improve the biological realism. Perhaps most importantly, we consider here the extreme case of the interaction becoming beneficial only in close proximity; one can also construct a model where interaction benefits are relevant even outside the collective. This would add Lotka-Volterra-type cross terms to the ecological dynamics captured by Eq. (1). Additionally, we set the collective’s carrying capacity *K*_*C*_ to a constant value independent of host or symbiont traits, since we envision that the mechanism of benefit exchange underlying this endosymbiosis enables the colonisation of a new niche (broadly construed) and that this niche has an associated carrying capacity. Better or worse usage of the niche (via different trait values) does not guarantee a higher maximum occupancy, only a quicker rate of reaching it. Further, we consider that the collectives must always be formed by a fixed density of symbionts acquired together and only once, i.e. we do not consider symbiont growth inside a single host. The size of these symbiont blocks does not affect the dynamics of our model per se, and so one can think of a single symbiont inside each host without loss of generality. However, in reality, each host has a dynamically changing symbiont population, and successive symbiont acquisition events would allow, e.g. decreasing symbiont population sizes to be propped up by immigration, or to be qualitatively changed by mutualism or antagonism between different symbiont strains (see (29)). Selection on the host due to the interactions with multiple symbionts might add further selection pressures that lead to a transition. Lastly, we also do not consider that there might be competition between the independent types and the collective due to niche overlap. Relaxing each of these assumptions is a worthy and important direction for future work.

**Table 3.**
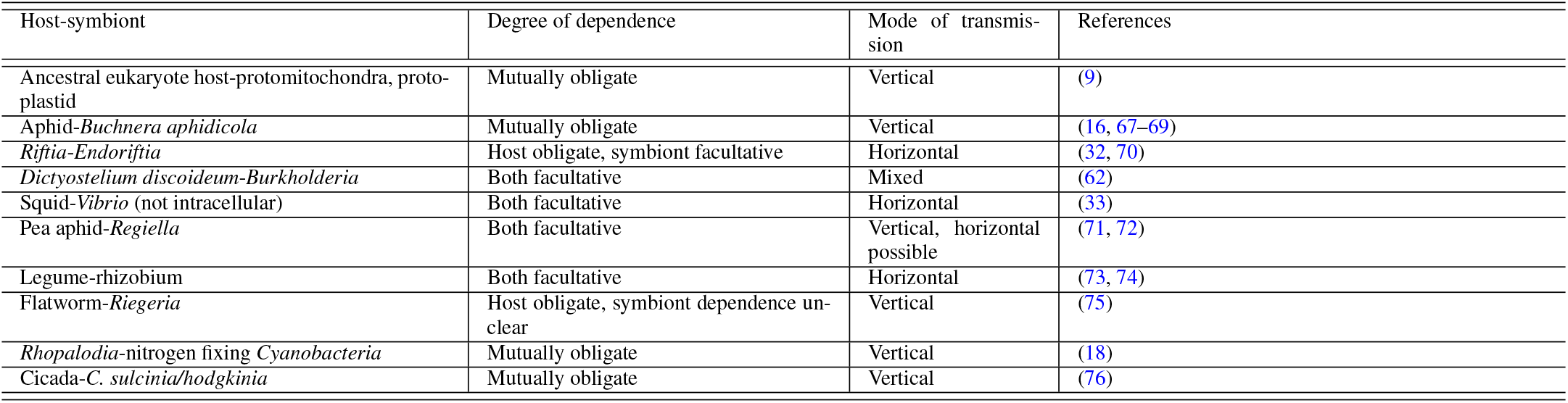
Dependence and cohesion information in some well-studied symbioses.

## Statement of Authorship

Conceptualization: CSG, PC, GSA; Funding acquisition: CSG; Model development and analysis: GSA, PC, CSG; Computer scripts: GSA; Supervision: CSG; Writing (original draft): GSA; Writing (review & editing): PC, CSG, GSA.

## Data and Code Availability

All data and simulation codes for generating the figures are available on Zenodo (https://doi.org/10.5281/zenodo.15533092).

## ACKNOWLEDGEMENTS

GSA thanks the DST-INSPIRE programme of the Government of India for financial support and the Max Planck Society for supporting the research visit. CSG and GSA thank the Max Planck Society for generous funding. Ananda Shikhara Bhat and Vishwesha Guttal helped with the translation of the abstract. We are grateful to Erol Akçay and two anonymous reviewers for useful comments.

This table is a survey of symbioses for which there is nontrivial information of the degree of dependence and the mode of symbiont transmission (a proxy for reproductive cohesion), adapted from a similar table compiled by (25). The information present, however, is (i) qualitative, (ii) a snapshot of the current state. The current state does not carry information about the full evolutionary trajectory, and hence is insufficient to estimate the traits Ω_*i*_ and *α*_*i*_. Therefore the available information does not allow us to confirm/refute the bias towards mutual dependence that our work predicts.

## Supplementary Note S1: Mathematical details of the model

### A. Traits and their relation to ecological parameters

Here we make explicit the interpretations of the traits Ω_*i*_ and *α*_*i*_. Recall that the obligacies Ω_*i*_ formalise a tradeoff between individual and collective reproduction. Further, the obligacy of an independent host does not affect the growth rate of the independent symbiont; the two traits only interact through their impact on the growth rate of the collective. Hence, if the growth rates of the three types are *f*_*H*_, *f*_*S*_, *f*_*C*_, then the corresponding constraints are

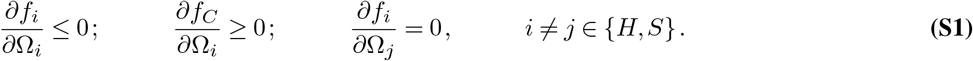

Therefore the intrinsic growth rate *f*_*i*_ of the population of species *i* is a function only of its own obligacy Ω_*i*_. Lastly, neither the host nor the symbiont can reproduce independently when its obligacy is 1, so *f*_*i*_(Ω_*i*_ = 1) = 0, *i ∈* {*H, S*}, and the collective cannot reproduce as a unit whenever any one of the host or symbiont are at zero obligacy: *f*_*C*_(Ω_*H*_, 0) = *f*_*C*_(0, Ω_*S*_) = 0 ∀ Ω_*H*_, Ω_*S*_ *∈* [0, 1]. We assume that *f*_*i*_ *≥*0 ∀*i* ∈ {*H, S, C*}.

The adhesions *α*_*i*_ formalise a tradeoff between synchronised and asynchronised reproduction. Therefore a higher adhesion increases the rate of association, increases the rate of collective reproduction, and decreases the rate of dissociation. If *a* is the rate of association and *d* is the rate of dissociation, we impose the constraints

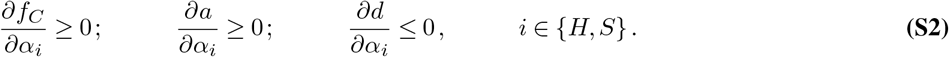

As rates, *a, d ≥* 0 at all trait values, and we assume that at *α*_*H*_ = *α*_*S*_ = 0, *a* is minimum and *d* is maximum; conversely at *α*_*H*_ = *α*_*S*_ = 1, *a* is maximum and *d* is minimum.

An example of equations that satisfy all these constraints is the following, and is the one (or variations thereof) we use across the main text:

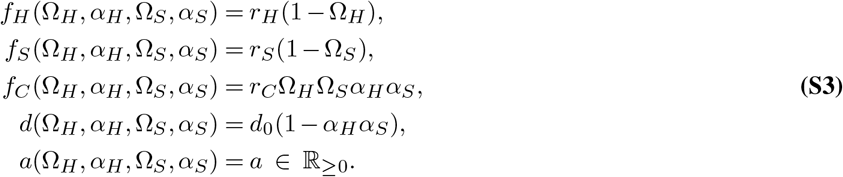

When the adhesions are held constant, this can be described by

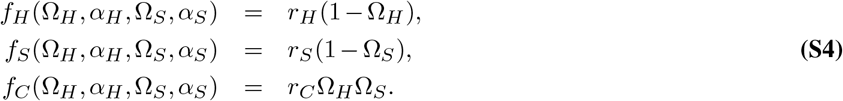

and when we assume obligacies to be constant, we model the ecological consequences of a different adhesion via:

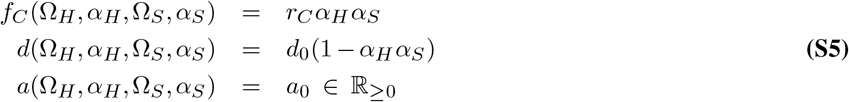

### B. Correspondence to a symmetric mutualism model

It is useful to establish the ecological context of our model and understand exactly how the presence of a collective affects the independent types. One can do this by comparing an extreme version (colloquially, “take a limit”) of our exponential model (Eq. system 8 in the main text) to more familiar models in the following way. Recall the population dynamics in the exponential model:

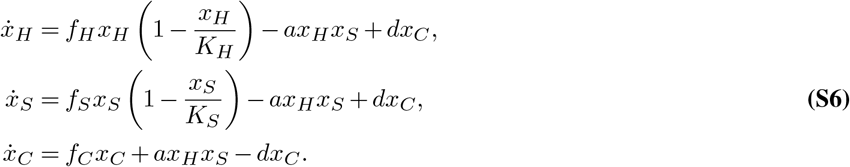

In all of our analyses, we are only interested in long-term symbioses like the nutrient-exchange symbioses in anaerobic ciliates (21), and not symbioses where the interaction is short-lived, like the parasite-cleaning mutualisms (77). If the interaction between the participants is ephemeral, the collective does not have as much of an independent existence. In the limiting case of instantaneous interaction, we can set 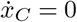. This has the effect of reducing the system of population dynamics equations Eq. (S6) to

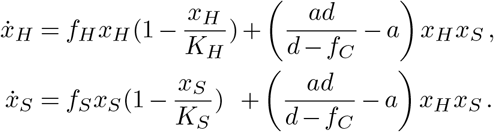

It is easily read off from this set of equations that there is now a constant *k* preceding the cross-term *x*_*H*_ *x*_*S*_ in both equations, with

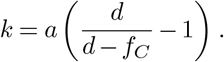

Since *f*_*C*_ *>* 0 by assumption, *d > d* − *f*_*C*_ and therefore *k >* 0 as long as *d > f*_*C*_. This last condition implies that the interaction is beneficial as long as *d > f*_*C*_ and becomes detrimental to both species when the collective growth rate *f*_*C*_ is “too high” - larger than *d*. Reinterpreting this cross-term *kx*_*H*_ *x*_*S*_ as arising from a Lotka-Volterra-type interaction, one arrives at the conclusion that the formation of a collective, in this “limit”, is equivalent to a symmetric mutualism as long as the dissociation is strong enough. If it is too weak, it goes from being a mutualism to being an interaction that is negatively affecting the growth of both parties. That it is symmetric in either case reflects the fact that with every collective reproduction event, there is exactly one new host and one new symbiont. This shows that collective reproduction can indeed be thought of as aligning reproductive interests of the host and symbiont. Like in a mutualism, the investment of one of the types (*H* or *S*) in collective reproduction therefore helps the other.

## Supplementary Note S2: Existence, dynamical stability, and variation of equilibrium population sizes

### A. Logistic model

The following figures show the variation of equilibrium population sizes 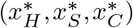 as a function of the obligacies (Ω_*H*_, Ω_*S*_) (Fig. S1) and the adhesions (*α*_*H*_, *α*_*S*_) (Fig. S2). All figures are in the form of heatmaps, and are obtained by Forward Euler-numerical integration of the population dynamical equations

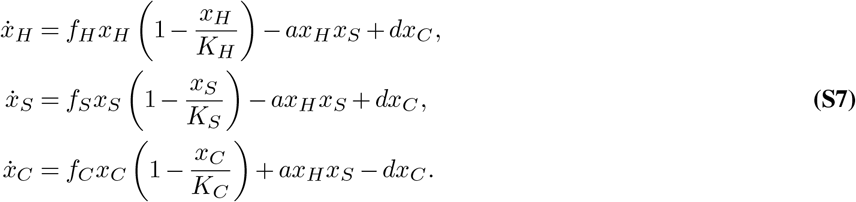

The above system is Eq. 1 in the main text. It is useful to note first of all that the routine found non-negative fixed points, and so one can say that biologically meaningful equilibria exist over a large range of parameters (over all values of traits, and over different values of *r*_*C*_, *K*_*C*_). Further, since the below equilibria were found by numerical integration, they are all necessarily stable (in the sense of a fixed point of a dynamical system). Third, the gradual change of the colours in the heatmap as the traits change suggests that it is in reality a single fixed point that changes location based on the trait values.

### B. Exponential model

The exponential model of population dynamics (Eq. (S6)) has four fixed points for the population densities, three of which are trivial. These trivial fixed points are located on the boundary at (0, 0, 0), (*K*_*H*_, 0, 0) and (0, *K*_*S*_, 0). The internal fixed point is given by

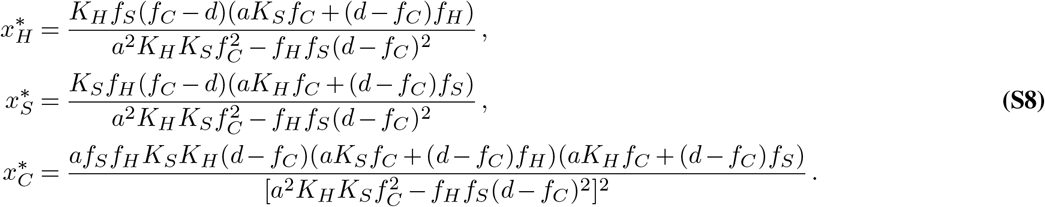

The interal fixed point 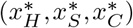 is internal, and feasible, i.e., all equilibrium densities are non-negative when

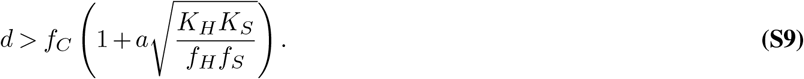

**Fig. S1.**
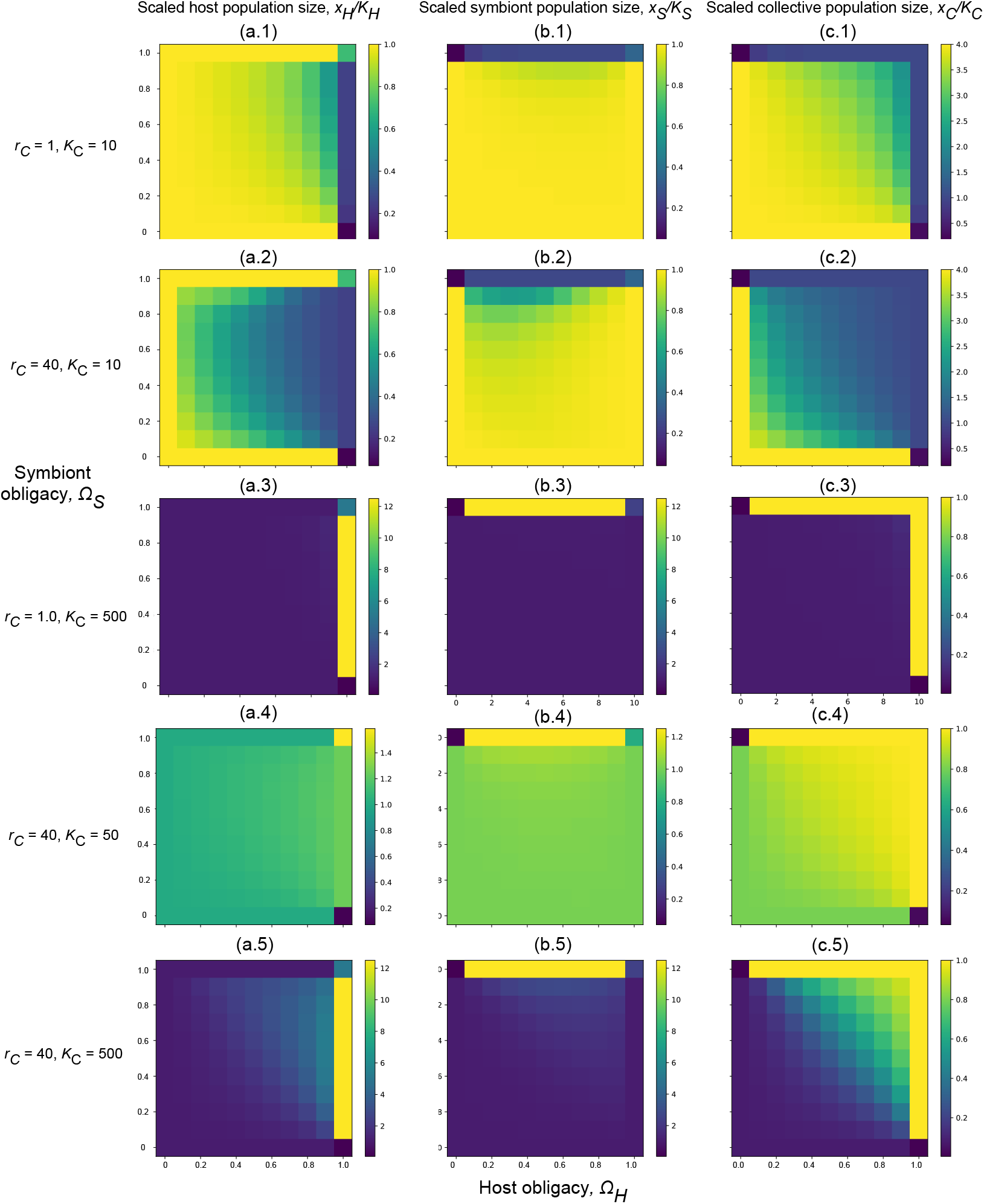
These panels contain heatmaps of the scaled equilibrium population sizes 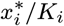 as functions of the obligacies Ω*i*, for five combinations of *rC* and *KC*. The first column (panels a.1 to a.5) corresponds to host population size, panels b.1 to b.5 to symbiont population size, and the last column to collective population size. The data for a given cell in a given heatmap is obtained by running a numerical integration routine (forward Euler) of Eq. (S7) starting from the initial conditions (*xH, xS, xC*)(0) = (10, 10, 0). The mapping from the traits to parameters is given by *fH* = *rH* (1 − Ω*H*), *fS* = *rS* (1 − Ω*S*), *fC* = *rC* Ω*H* Ω*S*. **Parameter values.** *rH* = 8, *rS* = 20, *KH* = 100, *KS* = 200, *a* = 0.1, *d* = 50.

To establish the condition for feasibility, first notice that any fixed point must satisfy

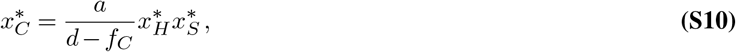

which arises from setting the dynamical equation for *x*_*C*_ to zero. First, notice that neither 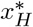 nor 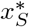 can be negative since we are searching for feasible equilibria. Therefore, we require *d > f*_*C*_ since otherwise 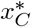 cannot be positive. The expressions for the host and symbiont population size have a similar form - they are, after all, symmetric in their attributes. Notice that the numerators of 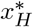 and 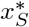 are negative whenever *d > f*_*C*_. Therefore, the expressions 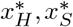 on the whole are positive whenever the denominator is also (strictly) negative. Then notice that they have the same denominator. The negativity of the denominator, upon rearranging to solve for *d*, exactly gives rise to the feasibility bound. Further, we determined computationally that this inequality guarantees linear stability for a large range of parameters (Fig. S3). Note that this lower bound on *d* is strictly larger than *f*_*C*_.

**Fig. S2.**
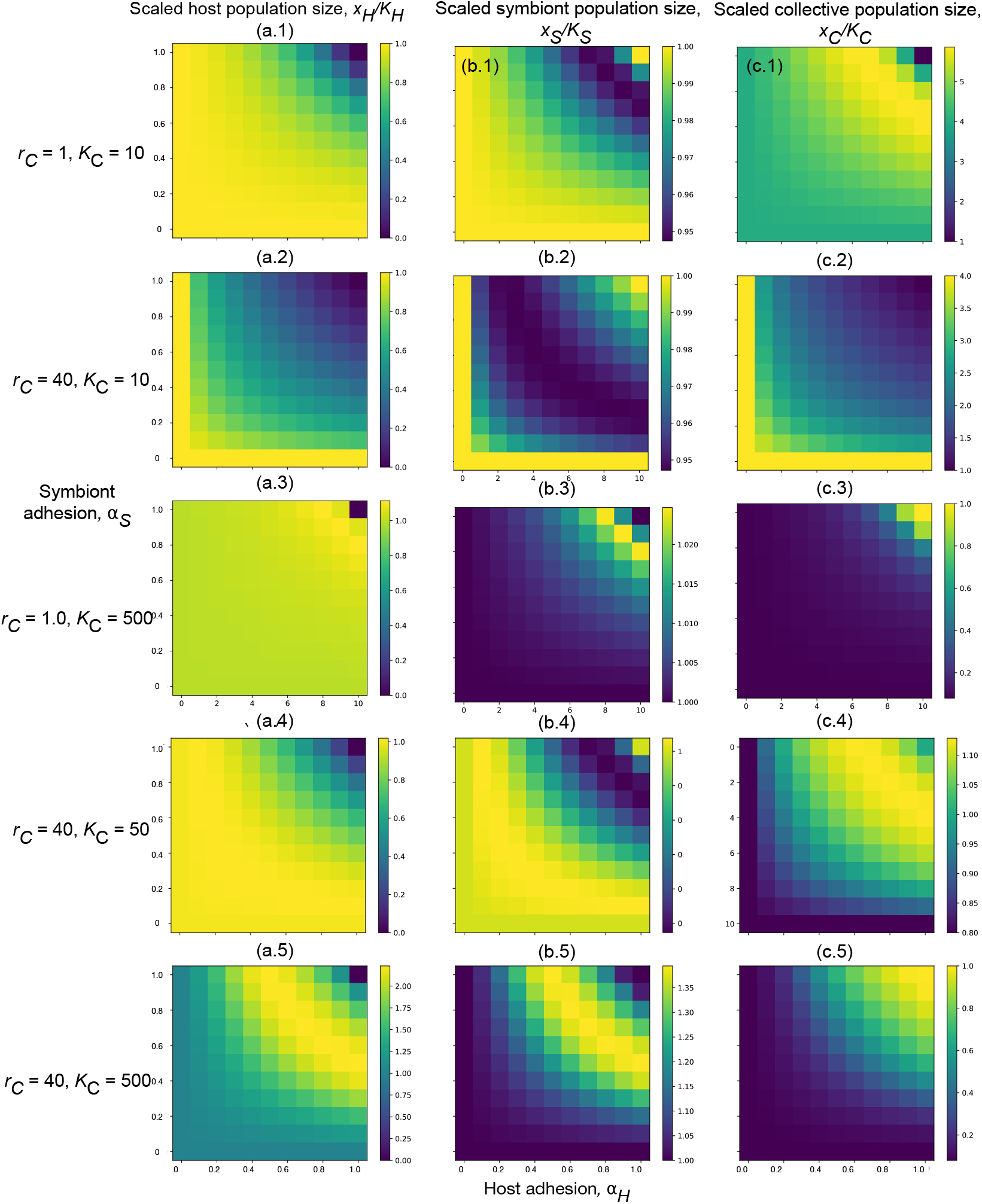
These panels contain heatmaps of the scaled equilibrium population sizes 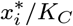 as functions of the adhesions *αi*, for five combinations of *rC* and *KC*. The first column (panels a.1 to a.4) corresponds to host population size, panels b.1 to b.4 to symbiont population size, and the last column to collective population size. The data for a given cell in a given heatmap is obtained by running a numerical integration routine (forward Euler) of Eq. (S7) starting from the initial conditions (*xH, xS, xC*)(0) = (10, 10, 0). The mapping from the traits to parameters is given by 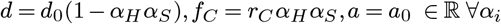. **Other parameter values.** *d*0 = 50, *a* = 0.1, *fH* = 8, *fS* = 20, *KH* = 100, *KS* = 200.

The next step is to find conditions for the stability of the equilibrium. The Jacobian of this system is given by

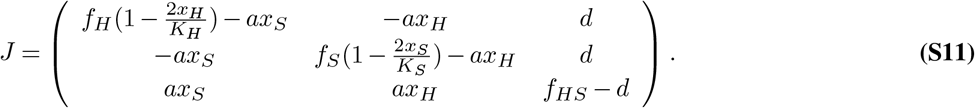

In the case of the trivial fixed point (0, 0, 0), the Jacobian above becomes upper triangular. The eigenvalues of a diagonal matrix are exactly the elements on its diagonal, and two of these elements are *f*_*H*_ and *f*_*S*_. We assume that these quantities are always positive in the entirety of this work, so this fixed point is never stable. It is not as easy for the other fixed points: the roots computed directly from the general characteristic equation are required and difficult to analyse, and we therefore resort to the Routh-Hurwitz criteria (78). This is a set of criteria on the coefficients in the characteristic equation of *J*. Specifically, let the characteristic equation be given by

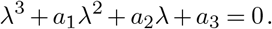

These numbers *a*_*i*_ are computable directly from *J*, and some of them are commonly known. For example, *a*_1_ above is the negative of the trace of *J*, and *a*_3_ is the negative of its determinant. The Routh-Hurwitz criteria state that all eigenvalues of *J*, i.e., all roots of the above polynomial, have a negative real part if and only if

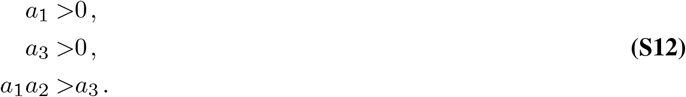

We shall not give the expressions for these coefficients *a*_*i*_ here, they can be found in the associated Mathematica notebook (‘./Analytics/obligacy-evolution.nb’).

For the two one-species equilibria (*K*_*H*_, 0, 0) and (0, *K*_*S*_, 0), the criteria can be verified easily: as long as the interior fixed point is feasible, these equilibria are unstable - one of the criteria above Eq. (S12) fails since *d > f*_*C*_. In particular, the feasibility bound being satisfied implies that *a*_3_ *<* 0 in these two cases. Further, there is a parameter range where none of the fixed points are stable: this is exactly when

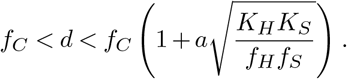

This is when the feasibility bound is violated (which is the right side of the above chain of inequalities), but not as strongly as to make *f*_*C*_ *> d* (left side of the chain). Therefore, the internal fixed point is rendered unstable, as well as all other fixed points: the only outcome for the abundances can be to escape to infinity.

For the internal equilibrium, the picture is much more involved. We cannot say much from the coefficients *a*_*i*_ themselves, but it can be shown that the subcriterion *a*_3_ *>* 0 exactly corresponds, upon some rearrangement, to the inequality that we have termed the feasibility bound. This proves that feasibility is necessary for stability.

Further analytics is difficult, and we therefore resort to numerically computing the maximum real part of the eigenvalues of *J* across a biologically reasonable range of parameter space. The results of this exercise are shown in Figure S3. This shows that the internal fixed point of the exponential model is, in fact, stable in the region of parameter space that we are interested in, and therefore, we proceed with further analysis.

## Supplementary Note S3: Independent trait evolution under the logistic model

### A. Adhesion evolution when host and symbiont are identical

Figure S4 demonstrates that the difference in host and symbiont population parameters induces the difference in their respective adhesion values’ evolutionary outcomes. The biologically realistic case is for *r*_*S*_ and *K*_*S*_ to be higher than the host counterparts, and this leads to the long run outcome *α*_*H*_ *<* 1, *α*_*S*_ = 1. It is trivial to observe that this outcome is flipped if the host now has the higher *r, K* – host and symbiont are just exogenously imposed labels, and can be flipped so that the old symbiont is called the “host”. In this figure, we show that if *r*_*H*_ = *r*_*S*_, *K*_*H*_ = *K*_*S*_ then there is no asymmetry in the trajectories – both host and symbiont adhesions stop short of (1,1) because it is not advantageous to go downhill from the local maximum in 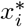 (as explained in the main text, using the support of Figure S2). Due to the stochastic nature of these simulations, some trajectories can escape to either side of the boundary; importantly, note that this happens in both host and symbiont trajectories.

### B. A sufficient condition for evolution to high Ω_*i*_ **or** *α*_*i*_

We claim that the following is a sufficient condition for the evolution of high obligacies or high adhesions:

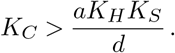

In particular, the right-hand side is the equilibrium population size of the collective that can be maintained by pure association and dissociation, i.e., how large the collective population is when *f*_*C*_ = 0, no synchronised reproduction. In Figure 4 of the main text, the parameter values are: *a* = 0.1, *d* = 50, *r*_*H*_ = 8, *r*_*S*_ = 20, *r*_*C*_ = 40, *K*_*H*_ = 100, *K*_*S*_ = 200. Therefore the value of 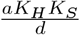 is 40. Our low value of *K*_*C*_ is 10, and the high value is 500, therefore satisfying the claim so far.

We tested the claim further by changing the value of *a* in two ways: first to decrease the value of *a* when *K*_*C*_ = 10 – this would push down the threshold 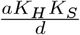 and make *K*_*C*_ high enough under our claim. The second test is to increase the value of *a* when *K*_*C*_ = 500, in an attempt to make *K*_*C*_ = 500 lower than the threshold. Both tests lead to results that support this criterion:

**Fig. S3.**
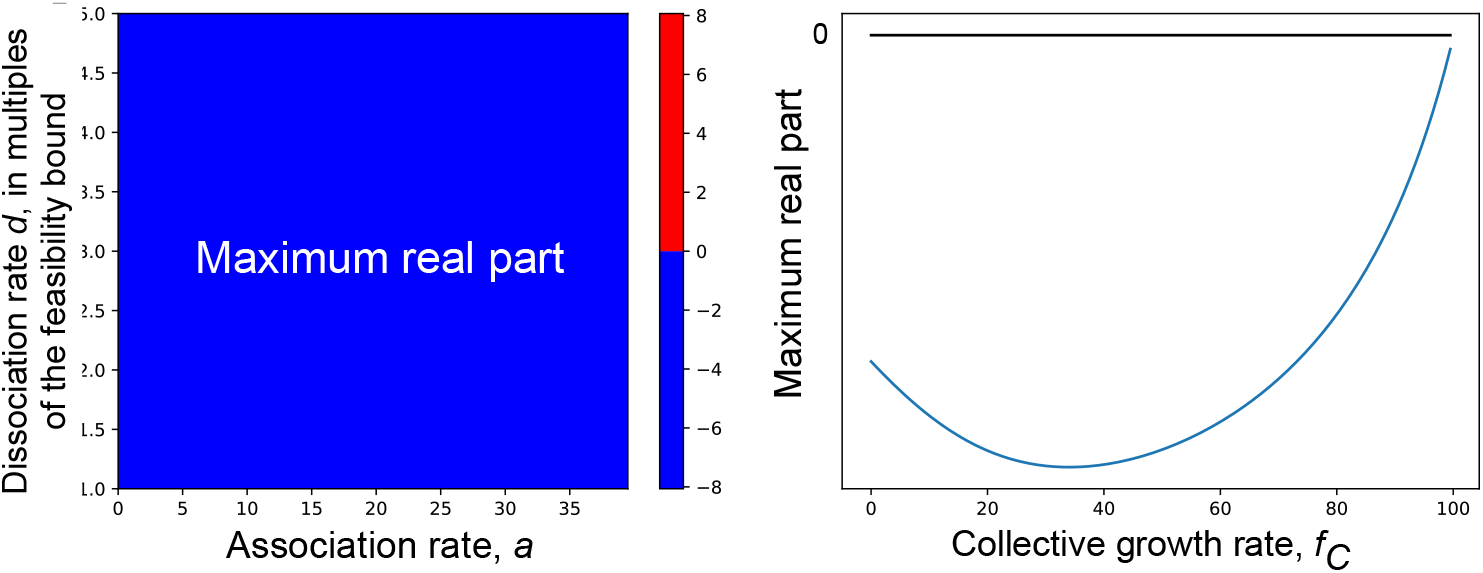
The internal fixed point of the exponential model is stable. Here we perform a numerical evaluation of the stability condition of the internal fixed point of the exponential model. In particular, we find and plot in two cases, the maximum real part of the eigenvalues of *J*, the Jacobian of the flow Eq. (S6). **Common parameter values for both panels**. *KH* = 100, *KS* = 200, *fH* = 8, *fS* = 20. **(a)** Red if unstable, blue if stable. Since we can already show that the feasibility bound is necessary for stability (see §B, we wish to only check stability when the feasibility bound is already satisfied. This panel is therefore a little unorthodox – the *x*-axis is the usual, but the *y*-axis is in multiples above the feasibility bound for the respective *x* value. Let 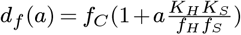 be the equalising value of *d* at the feasibility bound when all other parameters are kept fixed. Then in the heatmap, for a given value of the horizontal variable *a*, the vertical strip at this horizontal position ranges from [*df* (*a*), 5 *× df* (*a*)]. This panel therefore shows that the fixed point is stable for a large range of *d* values once the feasibility bound is crossed. **Parameter values**. *fC* = 10, *a ∈* [0, 40], *d ∈* [*df* (*a*), 5 *× df* (*a*)] **(b)** This shows that the fixed point is stable across a range of *fC* values. **Parameter values**. 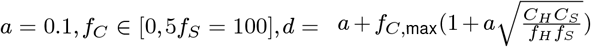 to maintain feasibility.

**Fig. S4.**
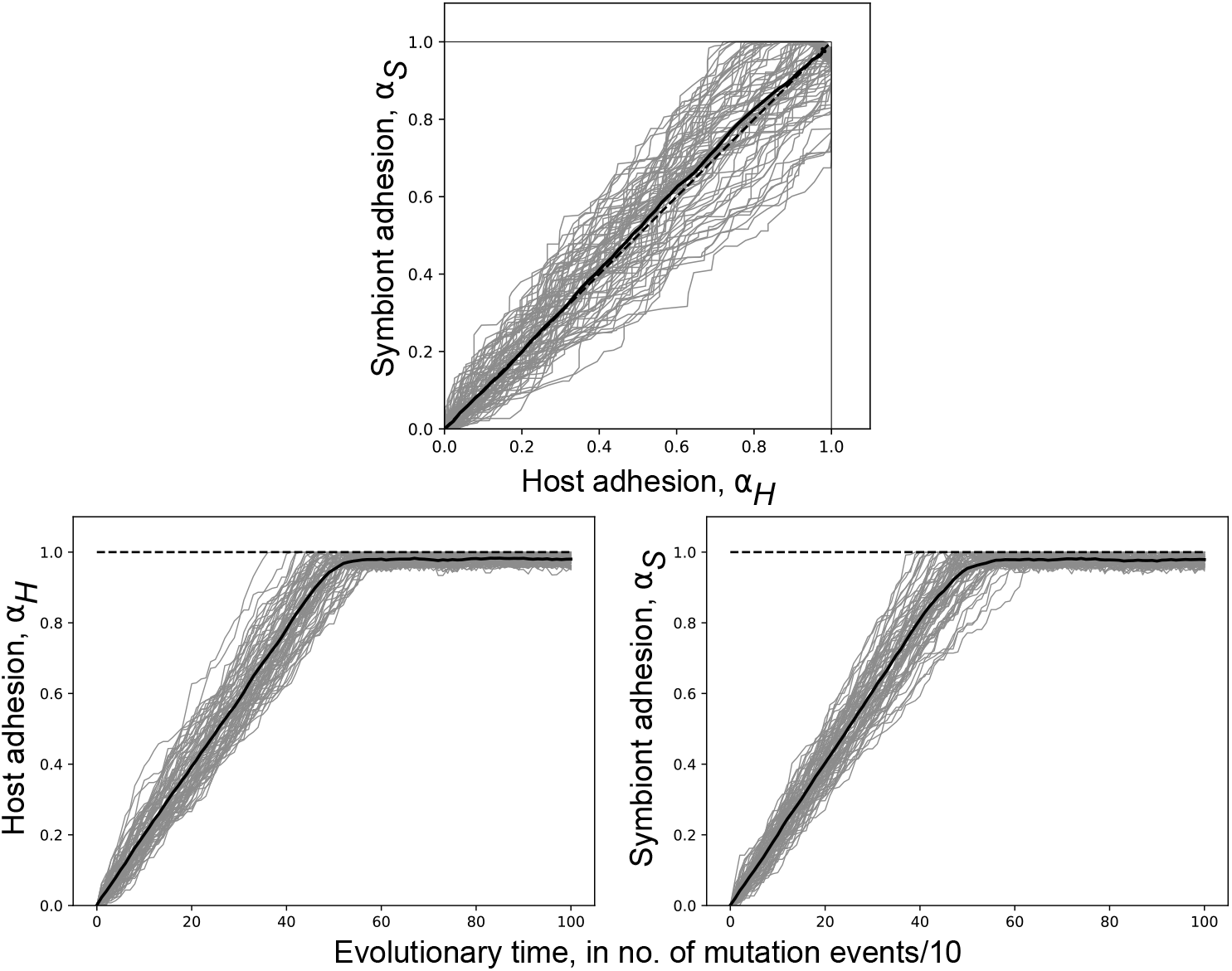
Adhesion evolution when host and symbiont are identical. Parameter values. *a* = 0.1, *d* = 50, *KH* = *KS* = 100, *KC* = 500, *rC* = 10, *rH* = *rS* = 8.

when *a* = 0.01, obligacies evolve to high values even when *K*_*C*_ = 10 (Fig. S5); when *a* = 5, obligacies remain at (0,0) even when *K*_*C*_ = 500.

**Fig. S5.**
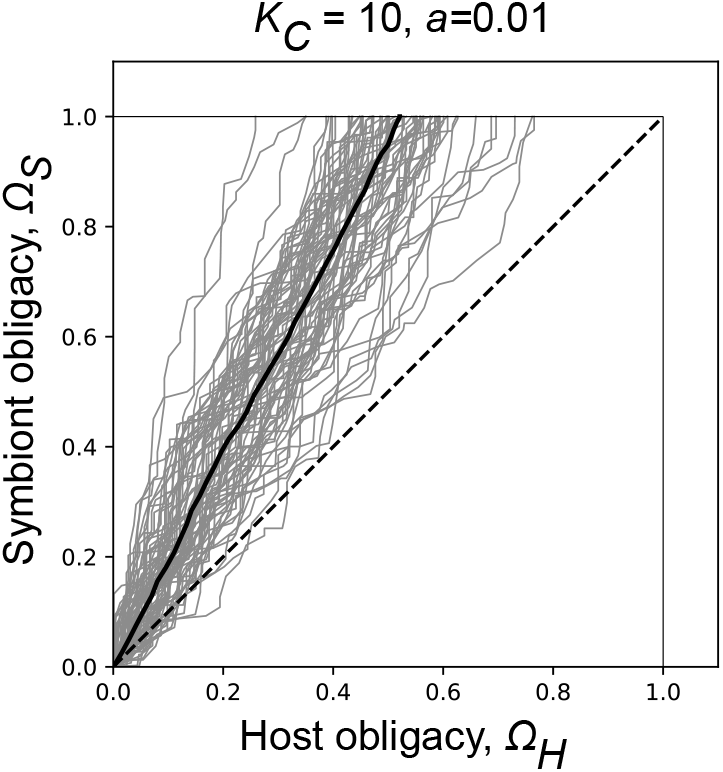
Parameter values. *d* = 50, *rC* = 40, *KH* = 100, *KS* = 200, 500, *rH* = 8, *rS* = 20.

However, due to a lack of analytical expressions, it is difficult to prove if this is also a necessary condition.

### C. Can (*α*_*H*_, *α*_*S*_) ever get to (1,1)?

Adhesion values can indeed reach (1,1) if *K*_*C*_ is high enough (see Figure S6). It can, of course, be argued that there is a point attractor at (1,1) that is only reached after infinite time and when *K*_*C*_ *→ ∞*, and that we only see it as (1,1) in these data due to the inherently discretised nature of simulations. However, this “one is only the supremum” interpretation is also good enough: the bottom line is that there is no upper bound on *α*_*H*_ that restricts it always to values less than 1.

**Fig. S6.**
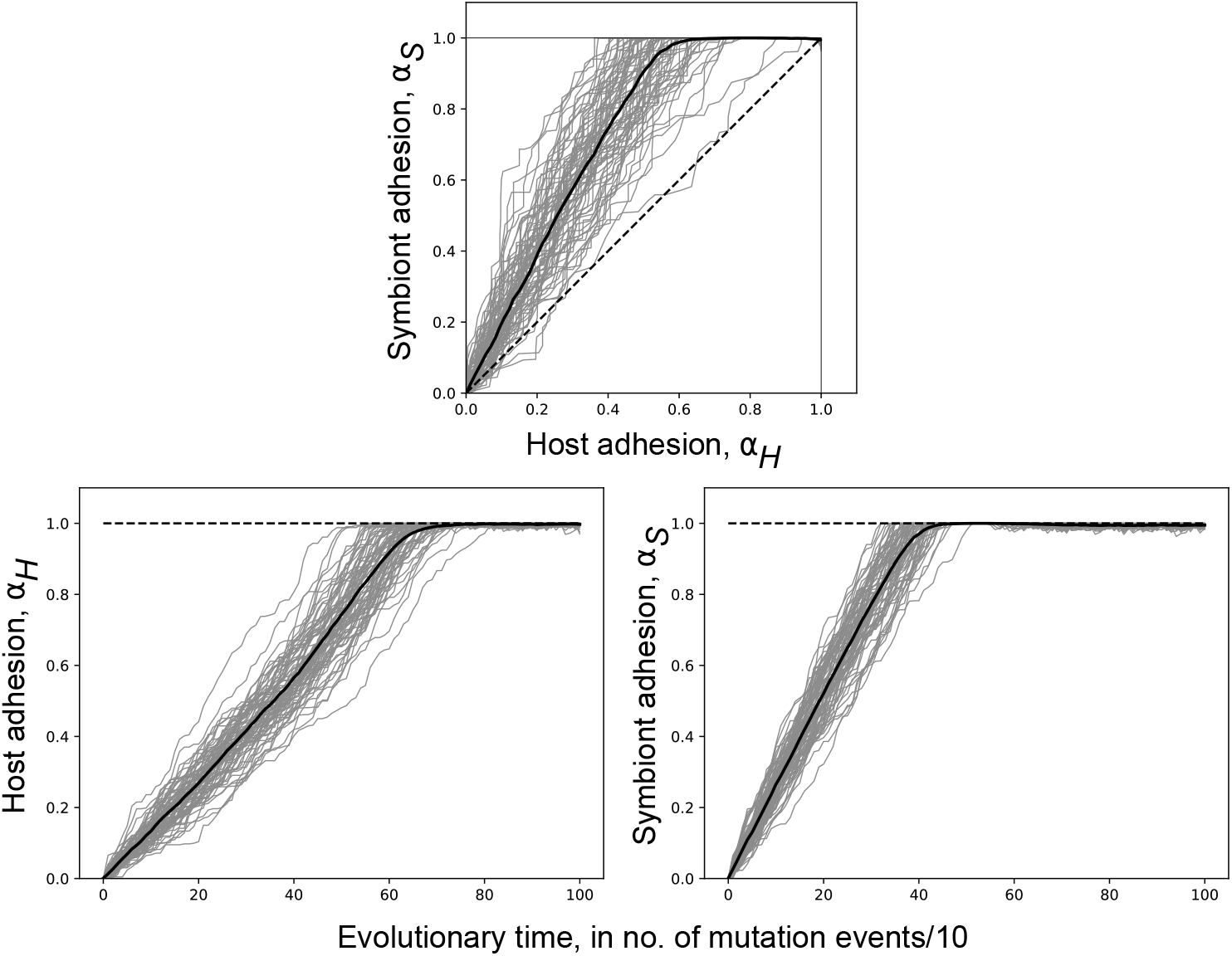
Parameter values. *a* = 0.1, *d* = 50, *KH* = *KS* = 100, *KC* = 10000, *rC* = 40, *fH* = 8, *fS* = 20.

### D. The effect of within-collective reproduction and mortality rates, additional figures

We reproduce the logistic model with within-collective birth and death rates (Eq. 6 in the main text) here for convenience:

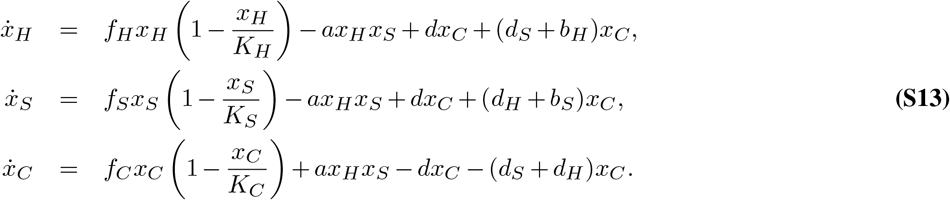

The effect of different values of *d*_*i*_ on the obligacies is presented in Fig. Eq. (S7), and the effect of different *b*_*i*_ values on obligacies is in Fig. Eq. (S9). The effect of different *b*_*i*_ and *d*_*i*_ on adhesions is presented in Fig. Eq. (S8).

**Fig. S7.**
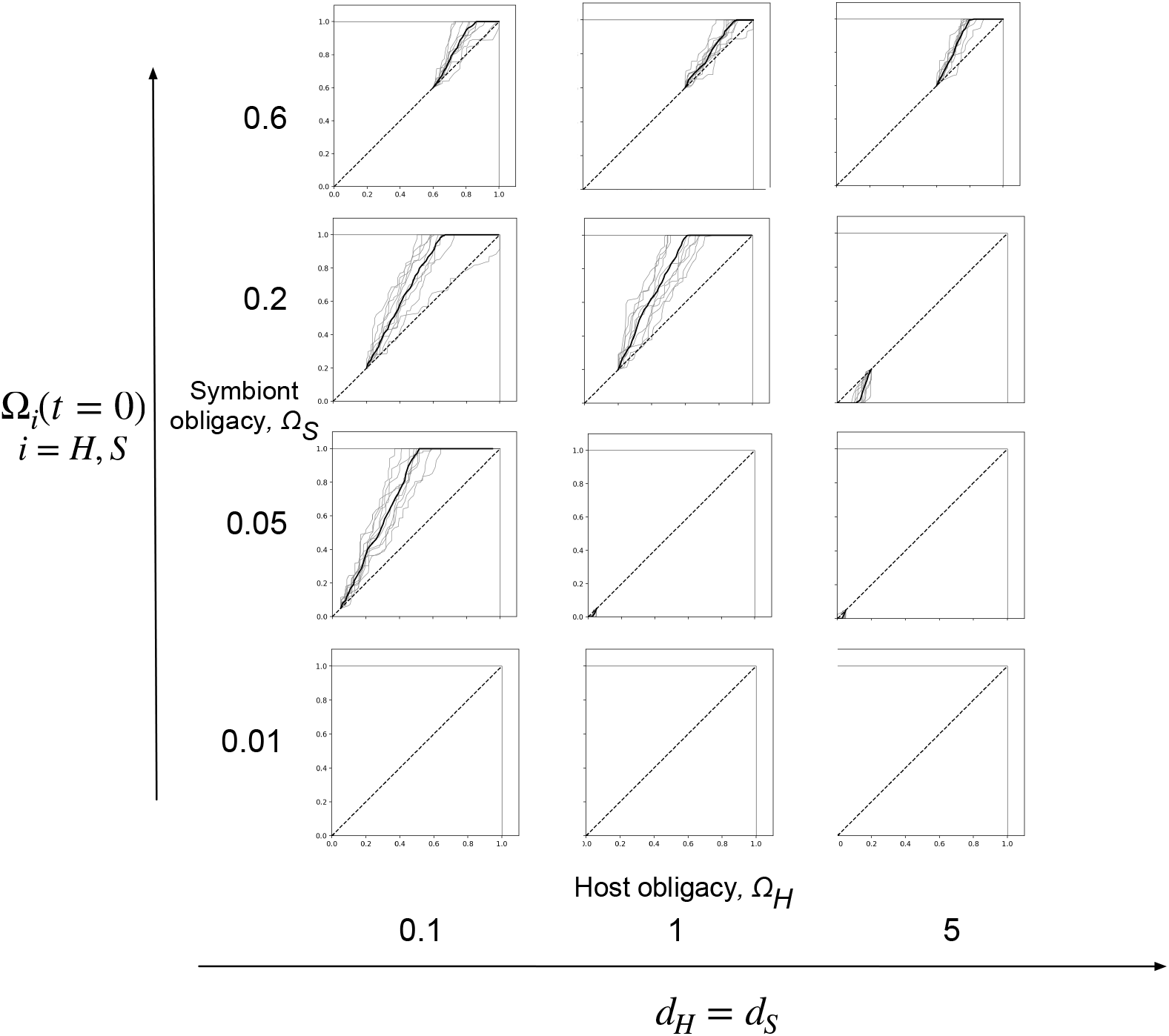
Adding a death term to obligacy evolution. **Parameters:** same as main text, *rH* = 8, *rS* = 20, *rC* = 10, *KH* = 100, *KS* = 200, *KC* = 250, *a* = 0.1, *d* = 50, *dH* = *dS ∈* {0.1, 1, 5}

## Supplementary Note S4: Deriving and using an invasion criterion

The population dynamical equations under the logistic model are given in the main text as Eq. system 1 and as Eq. S7 in this document.

To compute the invasion fitness of an arbitrary mutant, we suppose that there is a resident population with host and symbiont traits (Ω_*H*_, *α*_*H*_) and (Ω_*S*_, *α*_*S*_), respectively. After a sufficiently long time, a mutant arises when the resident population is at equilibrium, say with population sizes 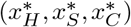. Due to the separation between ecological and evolutionary timescales, there can only be one mutant at a time, without loss of generality, say a host mutant. This mutant has trait value 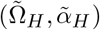 different from the resident, giving rise to different values of the parameters 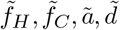 for the mutant host. The invasion condition is formalised as follows: we augment the model Eq. (S7) with additional equations tracking each type that arises due to the introduction of a mutant. Here, this always amounts to two more – the mutant host *y*_*H*_ itself and the mutant host-resident symbiont collective *y*_*C*_. The augmented model takes the form

**Fig. S8.**
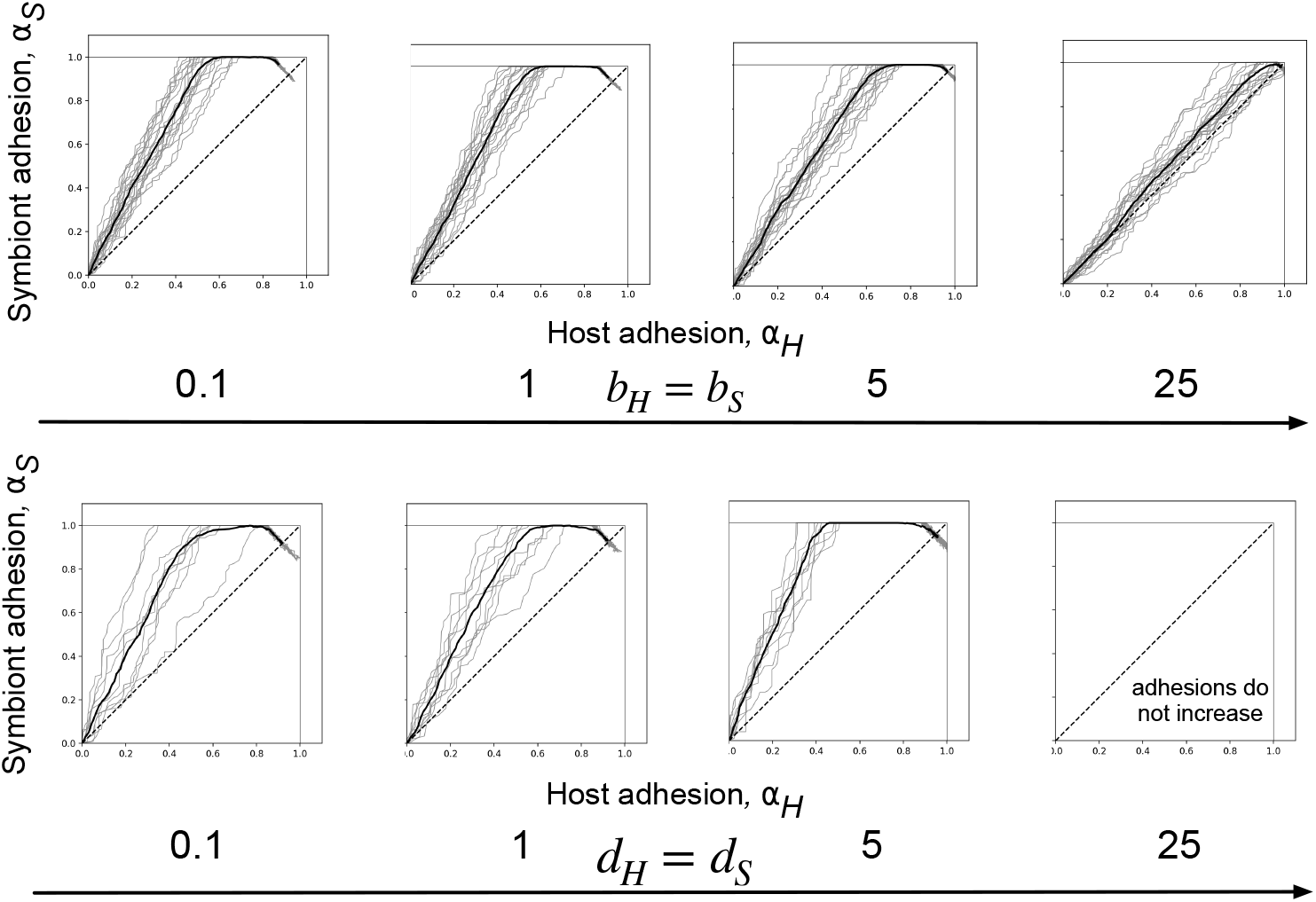
Adhesion evolution under within-collective birth and death terms. **Parameters:** same as main text, *fH* = 8, *fS* = 20, *rC* = 10, *KH* = 100, *KS* = 200, *KC* = 250, *a* = 0.1, *d*0 = 50. Top row: *dH*. = *dS ∈* {0.1, 1, 5, 25}, bottom row: *dH* = *dS ∈* {0.1, 1, 5, 25}

**Fig. S9.**
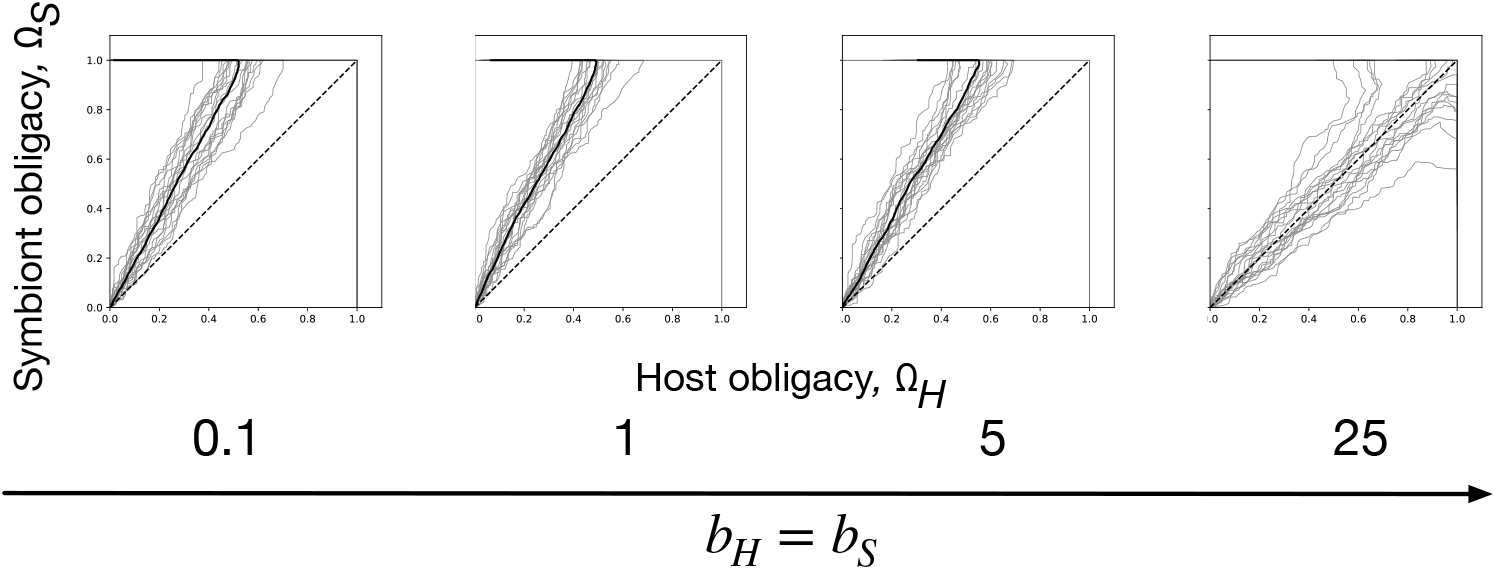
Obligacy evolution under within-collective birth term. **Parameters:** same as main text, *rH* = 8, *rS* = 20, *rC* = 10, *KH* = 100, *KS* = 200, *KC* = 250, *a* = 0.1, *d* = 50, *bH* = *bS ∈* {0.1, 1, 5, 25}

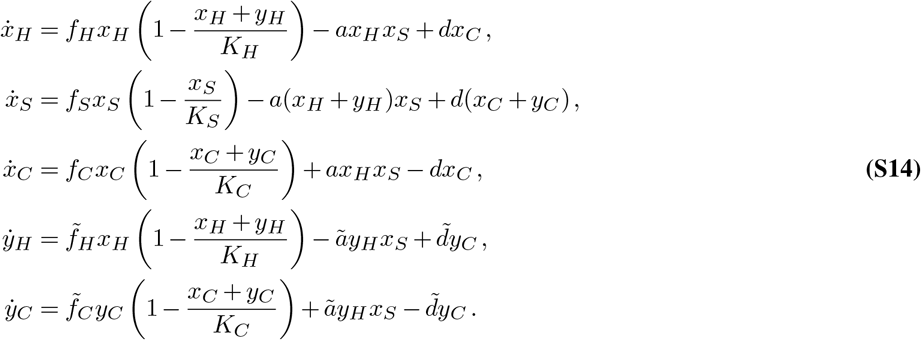

The invasion fitness is interpreted to be the growth rate of a rare mutant in a resident population that is at stable equilibrium (37). For a mutant host, this Jacobian (when the mutant is rare i.e., *y*_*H*_ = *y*_*C*_ = 0) takes the form

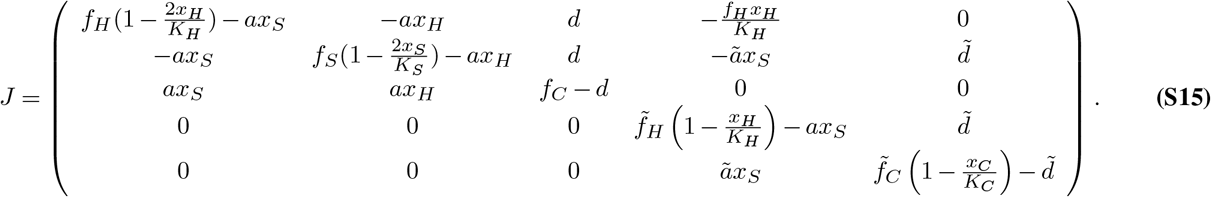

Due to the block-upper-triangular form of *J*, its eigenvalues are given entirely by the eigenvalues of only its diagonal blocks. We know the top-left 3 *×* 3 block has all eigenvalues with negative real part since we began with a stable population dynamical equilibrium. Therefore, we need only check the dominant eigenvalue of the submatrix corresponding to the mutant, i.e., the bottom right 2 *×* 2 block. The invasion fitness is thus defined as the dominant eigenvalue of the “sub-Jacobian” corresponding to the mutant equations, evaluated at 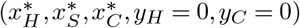:

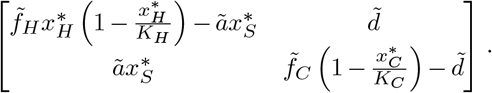

If it is positive, the equilibrium is destabilised in the presence of the mutant and the mutant can invade; if it is negative, the mutant goes to extinction (38, 46). This quantity is unwieldy and not amenable to mathematical analysis in our case, so we turn to other methods to quantify (in)stability of the resident population in response to mutants.

In particular, we use the next-generation theorem, which gives, under some conditions, an alternate characterisation of the standard stability condition of all eigenvalues having negative real parts (47, 48). Concretely, suppose we have a linear system of ODEs given by

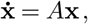

and the matrix *A* can be decomposed in terms of two matrices *F* and *V* as *A* = *F* − *V*, with some conditions on *F* and *V*. Now define *s*(*M*) to be the maximum real part of all the eigenvalues of a matrix *M*, and *ρ*(*M*) to be the maximum modulus of all the eigenvalues. The next-generation theorem (47) states that *s*(*A*) = *s*(*F* − *V*) has the same sign as *ρ*(*F*.*V* ^−1^) − 1. The quantity *ρ*(*F*.*V* ^−1^) is exactly the *R*_0_ value usually associated with the spread of virulent pathogens, and describes the average number of offspring one mutant individual sires. If it is larger than 1, then the mutant spreads. Therefore, if we can find a satisfactory decomposition *J* = *F*− *V* of *J* in Equation Eq. (S15), the fixed point in question is stable if and only if *ρ*(*F*.*V* ^−1^) − 1 *>* 0. This alternative method is useful because the latter quantity is sometimes more analytically tractable and interpretable than *s*(*J*).

### A. A useful but incomplete invasion criterion for the logistic model

For the logistic model, one can use the decomposition given by

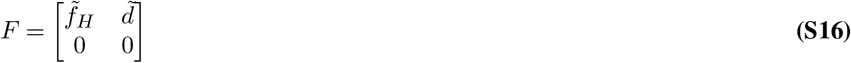

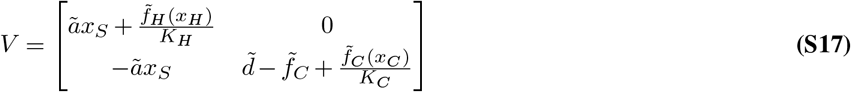

which satisfies all the requirements – all entries of *F* are positive, all eigenvalues of − *V* are negative (we assume that 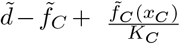 stays positive), and all elements of *V* ^−1^ are negative.

This leads to the invasion criterion

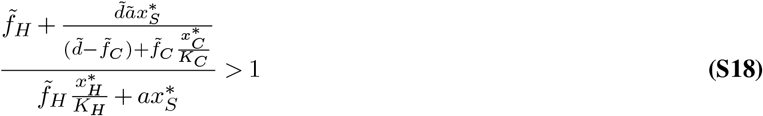

for a host mutant with traits 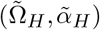 to invade a resident population composed of hosts having traits (Ω_*H*_, *α*_*H*_) and symbionts (Ω_*S*_, *α*_*S*_). Tilde-d quantities are functions of the tilde-d traits, and we assume *a* is independent of the trait values. Of course, a similar (but mirrored in H and S) expression exists for the invasion of a symbiont mutant. Although we know that the population sizes are themselves positive, it is difficult to solve exactly for when this quantity as a whole is positive because we do not possess expressions for 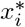 in terms of the other variables and parameters. However, we can use this to understand edge cases when we can indeed find the equilibrium population sizes analytically.

For example, one useful case is to consider Ω_*S*_ = 1, Ω_*H*_ *<* 1. Then *r*_*S*_ = 0, and the symbiont is sustained only by association and dissociation. This implies that at equilibrium, 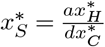. Further, since the association and dissociation must balance (see 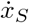 in Eq. (S7)), it follows that the host and collective populations must (each) be at either zero or their respective carrying capacities. Using this in *R*_0_ shows that it is exactly equal to 1, meaning that from here on, the value of Ω_*H*_ is not under selection.

### B. An exact invasion criterion for the exponential model

For the exponential model, we can use a similar decomposition

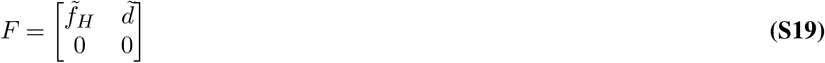

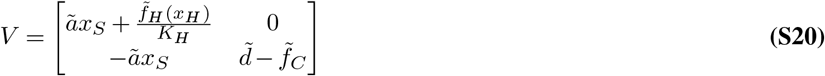

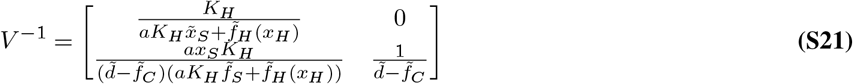

We also require that 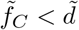 in the mutant host case, just as has been shown to be necessary for the existence of a feasible resident equilibrium. We now restrict ourselves to finding the condition under which *ρ*(*F*.*V* ^−1^) − 1 *>* 0. This leads directly to the invasion criterion

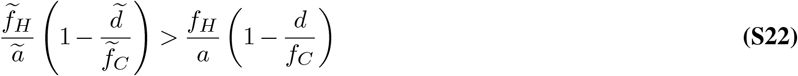

See the associated Mathematica notebook for more details.

The exponential model, however, has an associated artefact – the exponential growth of the collective destabilises the internal fixed point before (1,1), in contrast to the logistic model. In Fig. S10, we tracked the independent evolution of Ω_*i*_ (panels a,b) and *α*_*i*_ (panels c,d) in a particular system with realistic parameters while numerically checking if feasibility and stability hold throughout evolution. Consider the evolution of obligacies (and similarly for adhesions): Starting from (Ω_*H*_, Ω_*S*_) = (0, 0), i.e., no dependence, we observe first that both obligacies increase, as expected. However, the trajectory does not reach (Ω_*H*_, Ω_*S*_) = (1, 1). The obligacies increase to a certain point, making the ecological equilibrium unstable and infeasible. The feasibility bound Eq. (S9) is violated, leading to the collective growth rate becoming too high to sustain equilibrium host/symbiont populations. This barrier in the Ω_*H*_ − Ω_*S*_ plane is artificial due to the unrealistic assumption that the collective population can grow without bound. We therefore “trust” the results of the exponential model only when the traits have low enough values, and in this regime, we can take advantage of the analytics developed above.

Stepping away from the mathematical machinery, there is a clear explanation for the evolution of higher Ω_*i*_ and *α*_*i*_ - as a result of the collective’s exponential growth, its population grows without bound; so the investment of the host/symbiont in *f*_*C*_ is not unexpected. While this result is, therefore, not surprising in itself, the analytical setup is conducive to asking more nuanced questions.

### C. Long-term evolutionary dynamics in the exponential model via the canonical equation

Suppose, for simplicity, that the mutant has obligacy 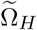 and the resident has obligacy Ω_*i*_. The invasion fitness is given by

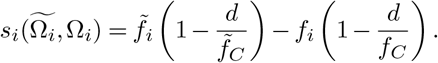

The invasion fitness determines the fate of a given mutant and, therefore, induces a certain path for the evolutionary dynamics of the trait value. This path is given by the canonical equation of adaptive dynamics (38), which is an ordinary differential equation describing the macroevolutionary change, and the obligacy of species *i* varies as

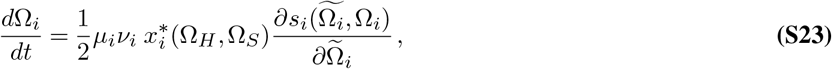

where *µ*_*i*_, ν_*i*_ are respectively the mutation rate and the variance of the mutation step, 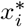 is the equilibrium population size, and the partial derivative is the fitness gradient. By the constraints Eq. (S1) and the fact that *d > f*_*C*_, one sees that the fitness gradient is uniformly positive.

A similar analysis can be accomplished when a mutant has both different obligacy and different adhesion values. Again, tilde-d quantities are associated with the mutant. The invasion fitness is given by

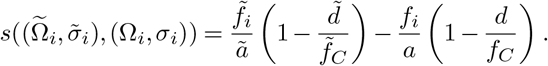

**Fig. S10.**
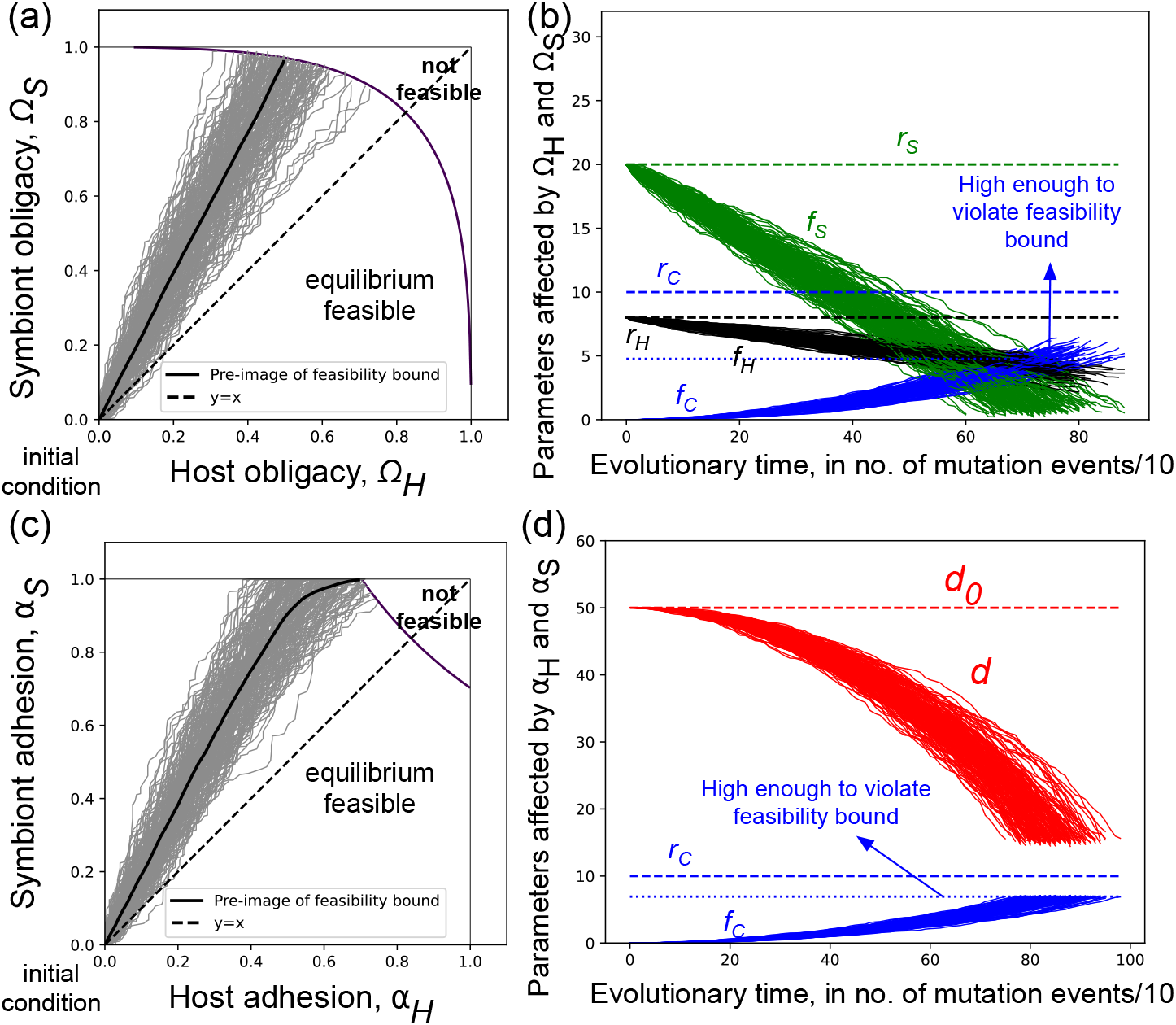
Evolutionary trajectories are biased towards higher symbiont obligacy. **(a**,**c)** For the mappings in Eqs. Eq. (S4) (panels a,b) and Eq. (S5) (panels c,d), we simulated 200 independent evolutionary trajectories. The trajectories monotonically increase with Ω*S* (a) and *αS* (c) increasing faster respectively, and then stop when the fixed point is no longer stable – this happens precisely at the feasibility bound Eq. (S9). In the infeasible region, the collective population density increases to infinity and, due to dissociation, maintains a comparatively negligible population of independent hosts and symbionts. (See SI §S.5 for information on calculating the average trajectory). **(b**,**d)** Using the mappings, we can also visualise the dynamics of the growth rates (in a) and the association-dissociation rates (in d). For example, in (b) the independent growth rates decrease, and the collective growth rate increases until it becomes high enough that the feasibility bound is violated. **Parameter values**. (a,b): *rH* = 8, *rS* = 20, *rC* = 10, *KH* = 100, *KS* = 200; for (c,d): *fH* = 8, *fS* = 30, *rC* = 10, *KH* = 100, *KS* = 200, *a*0 = 0.1, *d*0 = 50.

We write now the canonical equation for a multi (here, two)-dimensional trait:

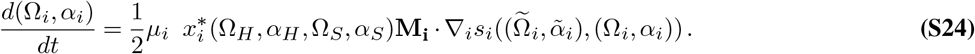

Here *µ*_*i*_ is the mutation rate for type 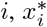 is its equilibrium population size at the given trait values, and **M**_**i**_ is the variance-covariance matrix of the traits of type *i* which encodes correlations between these traits. *s*_*i*_ is the invasion fitness for type *i*, and the gradient is taken with respect to all of its traits. This more general equation is similar in spirit to the earlier equation Eq. (S23), with changes accounting for the now arbitrary (in ℕ) dimension of the trait.

To show that the traits increase, it is sufficient to show that all the entries of 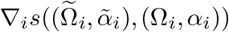 are positive. Under the constraints on the partial derivatives that we set up initially, and some additional similar constraints on the mapping from traits to ecological parameters, it is straightforward to show this.

### D. The impossibility of branching when the invasion fitness is separable

Let *s*(*y, x*) be the invasion fitness of a mutant with trait *y* in the background of a resident population of trait value *x*, with traits taking value in some set *T*. We say that *s*(·,·) is *separable* if there exists a function ℱ: *T →* ℝ _*≥*0_ such that one can write *s*(*y, x*) = ℱ (*y*) − ℱ (*x*).

Recall that a mutant invades when its invasion fitness is positive. Therefore, if the invasion fitness is separable, a mutant with trait *y* invades exactly when ℱ (*y*) *>* ℱ (*x*). We notice, then, that an invasion fitness is separable when there is some function ℱ (·) that acts like a fitness - if *y* has a higher value of ℱ, then it invades, and if it has a lower value of it ℱ goes to extinction. Evolutionary trajectories hence maximise this quantity, and therefore, in this sense, the fitness landscape does not change. If the invasion fitness is separable, one can construct a fitness landscape ℱ on the trait space such that all permitted trajectories climb up this landscape. Now we prove the observation that we made in the main text regarding the invasion criterion.

Suppose *s*(*y, x*) is the invasion fitness describing the evolutionary dynamics of a given system, and that it is separable. Then we will show below that a given strategy *x*^*\**^ is uninvadable if and only if it is convergence stable. First we define the function

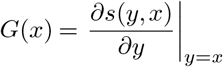

which is the fitness gradient for an invading mutant of trait value *y* against a resident with trait *x*. We shall use the fact that for any (*C*^2^) function,

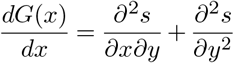

But since *s* is separable, there is a function ℱsuch that *s*(*y, x*) = ℱ (*y*) −ℱ (*x*). This implies that 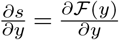since ℱ (*x*) does not depend on *y*, and then 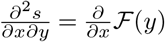 which is obviously zero. So we have shown that

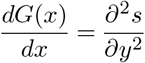

The left-hand side determines convergence stability, and the right-hand side determines (un)invadability, and since they are equal, these notions cannot diverge!

Recall that a singular strategy is called a branch point, i.e., a point at which evolutionary branching occurs, if it is convergence stable but invadable. A corollary of the above result, then, is that separable invasion fitnesses do not admit branch points - evolutionary branching is impossible. It is perhaps useful to think in terms of the contrapositive: if evolutionary branching is possible, then the invasion fitness is *not* separable.

This is because for a singular strategy to be a branch point, it must by definition be convergence stable but invadable by similar mutants (39). This is possible in general because the conditions to reach a maximum (convergence stability) and to stay there (invadability) are not the same. However, if one can identify a quantity that uniformly increases along all permissible evolutionary trajectories, these notions cannot be different. The reason why is simple and goes as follows. When such a quantity exists, uninvadable points are all convergence stable since they can be reached by trajectories that maximise this quantity. Conversely, convergence stable singular strategies are always uninvadable since any trajectory that converges there must maximise this quantity.

### E. Trajectories in the dependence-cohesion plane

In this section, we provide the supporting information for the result that mutual dependence evolves faster than reproductive cohesion. We will first provide an explanation, in terms of the model and then biologically, for why it is adaptive to invest more in obligacy than adhesion. Second, we will show that the bias towards mutual dependence is robust to other minor choices that we made in arriving there.

For providing an explanation, consider the trait-mapping that we have utilised in the main text (Eq. 2 there), also mentioned as Eq. (S3) in this document.

We claimed in the main text that since a change in *α*_*H*_ has the functional effect that depends also on *α*_*S*_ (and vice versa), it has a lesser adaptive benefit than a change in Ω_*i*_. To show this, notice that two important choices were made in arriving at the above trait mapping – first, that the independent growth of the host and symbiont depends only on their own obligacy; second, the dissociation rate depends on both *α*_*H*_ and *α*_*S*_. However, in general, the independent growth rates can depend on either (i) only their own obligacy or (ii) both obligacies; the dissociation rate can depend on (I) only one adhesion and (II) both adhesions. The “default” map that was motivated by biological reasoning corresponds to the combination i-II above. We shall below give the trait mapping of all other combinations, with each followed by the induced evolutionary trajectories of the traits Ω_*i*_ and *α*_*i*_. There is another note to be made here: when we make the association and dissociation rates dependent only on one of *α*_*H*_ and *α*_*S*_, we pick the symbiont trait without loss of generality. This does not have a qualitative effect, and the exact effect will be made clear when necessary. Further, we fix the effect of the obligacies and adhesions on *f*_*C*_; this does not change because our mental model of an endosymbiosis still stays the same - increasing both of these traits improves collective growth rate. The association rate *a* will remain a constant here, like in the rest of this work.

In all of the below cases, there will be some commonalities - all four traits always increase, and within each trait pair, the symbiont trait increases faster. This is due to the fact that the exponential growth of the collective and the difference in parameters are not changed over the course of this exercise. The traits increase due to the exponential growth of the collective and the fact that *f*_*C*_ can be increased by any one of the four traits. The symbiont trait increases faster than its corresponding host trait due to the different generation times and carrying capacities.

Now, consider the combination i-I, with results in Figure Eq. (S11). Here, the trait mapping takes the form

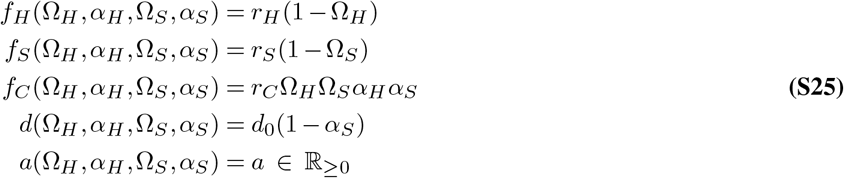

**Fig. S11.**
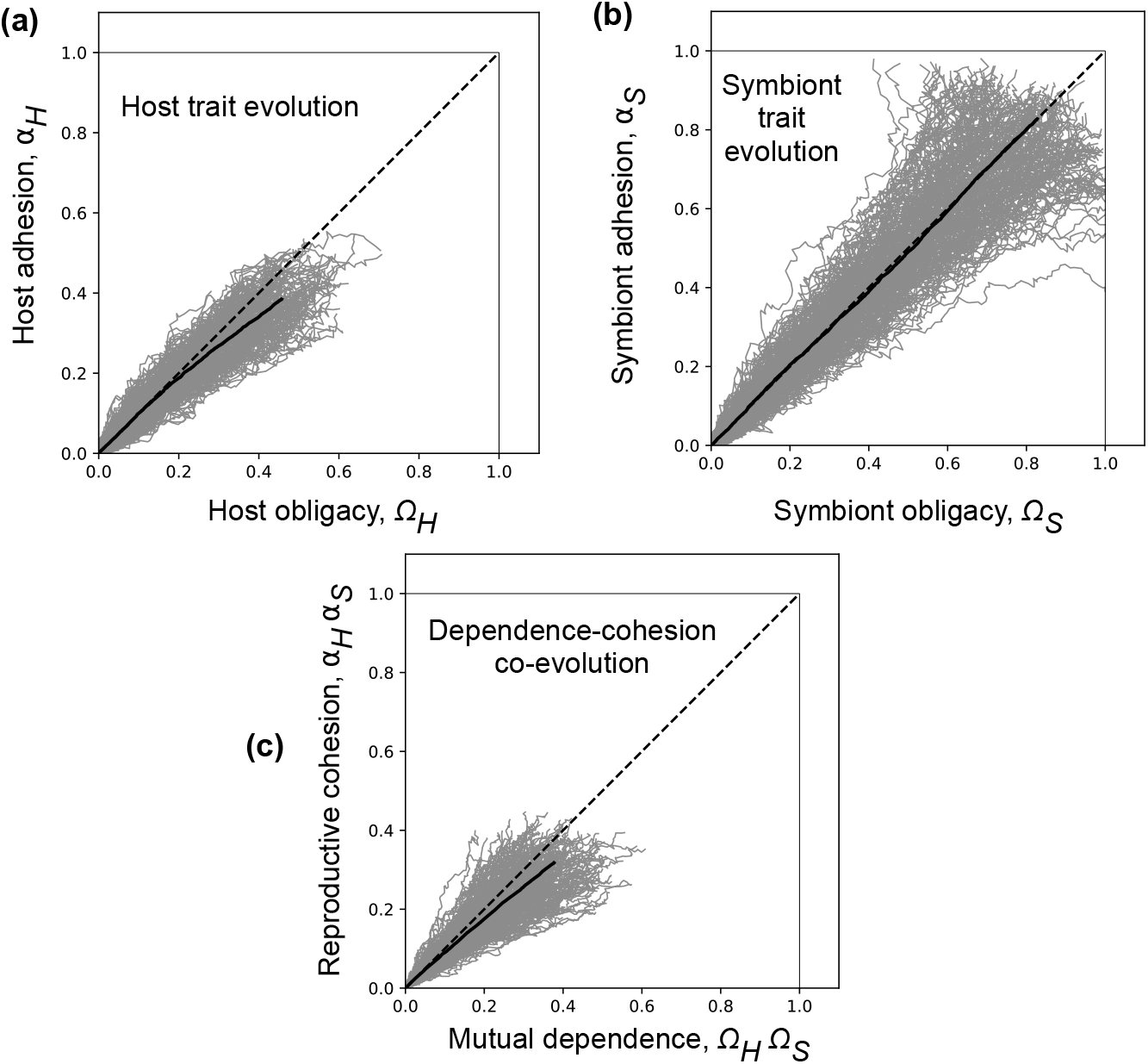
Independent growth rate depends only on own obligacy, dissociation rate depends only on one adhesion.

Here, there is a slight bias towards mutual dependence, which can be understood by looking at the evolution of the host and symbiont in their own trait space. There is a benefit for both host and symbiont to increase obligacy, and there is the *same* kind of benefit for the symbiont to increase its adhesion since *d* = *d*_0_(1 − *α*_*S*_). There is a smaller benefit for the host to increase its adhesion because an increased host adhesion only increases *f*_*C*_ but doesn’t decrease *d*. Therefore, Ω_*H*_ evolves faster than *α*_*H*_ ; however, Ω_*S*_ and *α*_*S*_ evolve at the same rate. If instead *d* = *d*_0_(1 − *α*_*H*_), there is no bias between Ω_*H*_ and *α*_*H*_ whereas Ω_*S*_ evolves faster than *α*_*S*_. This leads to a slight bias towards mutual dependence. This bias is a little artificial in the following sense: the bias arises only because we are forced to make a choice between *α*_*H*_ and *α*_*S*_ in the form of *d*, and would not exist if this choice were not thrust upon us.

Second, consider the combination ii-I with results in Figure Eq. (S12) and the following trait mapping:

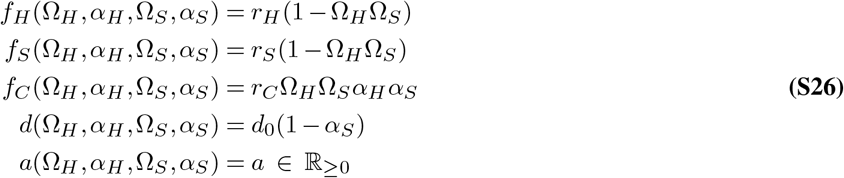

Here, there is a bias towards reproductive cohesion. This is because the benefit of increasing the adhesion is now stronger than that of increasing obligacy. This is further, to be more precise, because the functional effect of increasing adhesion also does not depend on the other adhesion. There is also a slight bias towards dependence in the host because increasing adhesion for the host does not increase d, but again this is because we are forced to make the same choice between *α*_*H*_ and *α*_*S*_ for *d*. If instead *d* = *d*_0_(1 − *α*_*H*_), there would be a slight bias towards dependence in the symbiont and a bias towards cohesion in the host.

**Fig. S12.**
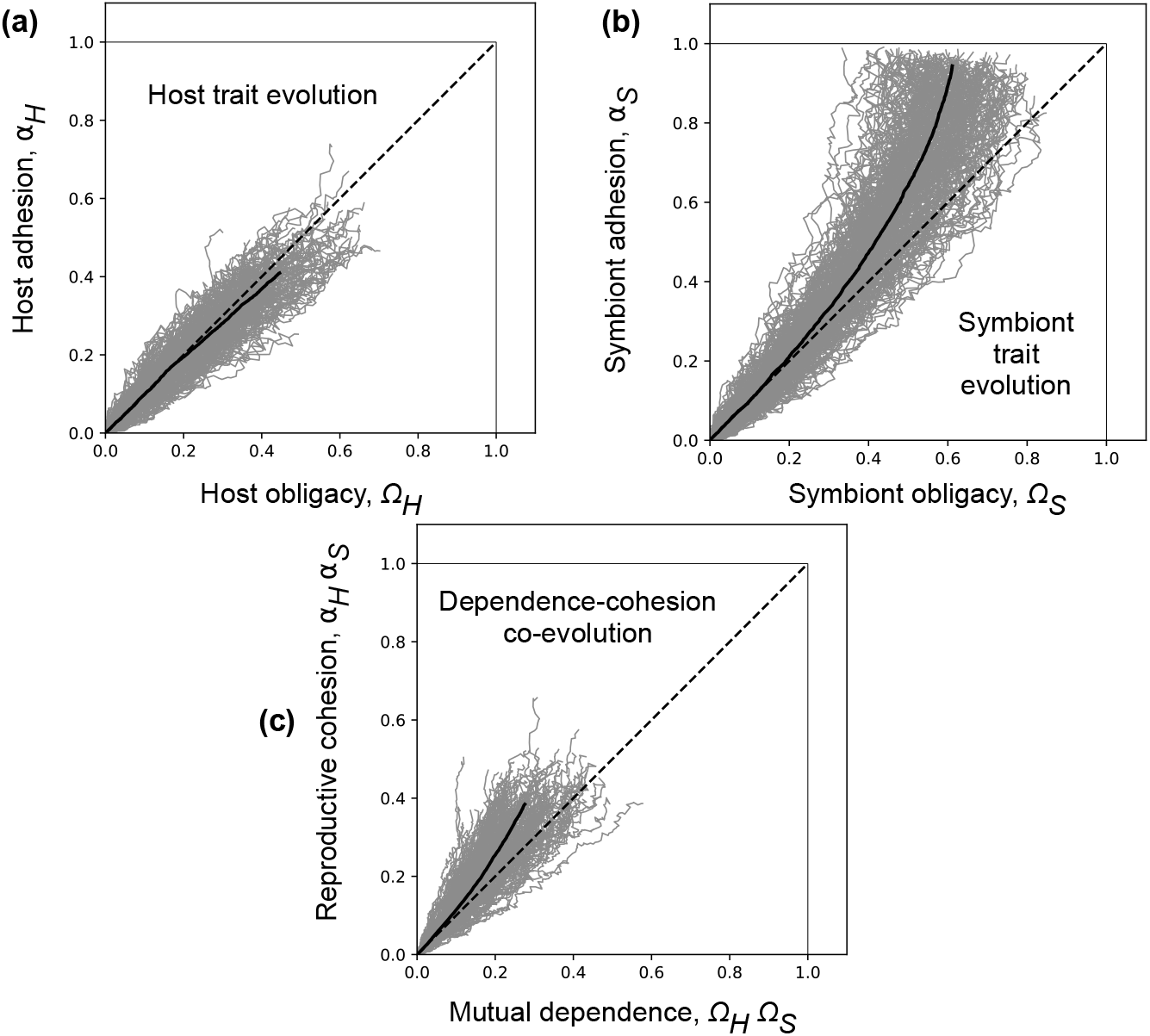
Independent growth rate depends on both obligacies, dissociation rate depends on only one adhesion.

**Table S1.** A schematic description of trajectories in the dependence-cohesion under different kinds of mappings, with the shorthand i-ii-I-II described in Section E. There is a pattern in this table: when both trait pairs Ω*i* and *αi* combine in the same way to influence ecological parameters (i-I or ii-II), there is little to no bias. The slight bias in i-I case is an artefact, see main text for an explanation. On the other hand, when they combine in different ways (i-II or ii-I), there is a bias towards faster evolution of dependence in one case and faster evolution of cohesion in the other. Our work arrived at case i-II from biological considerations, and we therefore make the associated prediction.

|  | i | ii |
| --- | --- | --- |
| I | Slight bias | Bias towards reproductive cohesion |
| II | Default, bias mutual dependence | No bias |

Recall that the default combination that we have worked through in the main text is given, in this shorthand, by i-II, so we will not consider it again. Now, lastly, let us study the combination ii-II with results in Figure Eq. (S13) and trait maps:

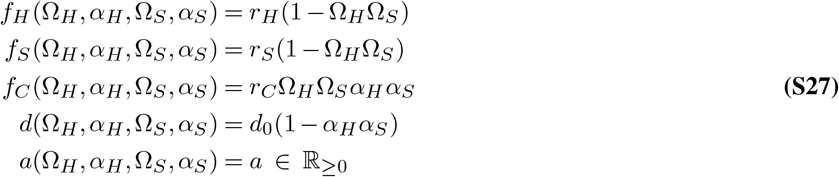

There is, as expected, no bias between mutual dependence and reproductive cohesion. Table S1 summarises the above results. Now, we shall show that the bias towards mutual dependence is robust to the choices that we made throughout the main text in arriving at this result. For ease of reference, we present again the alternate trait map relevant for panel **(a)** of Figure S14

**Fig. S13.**
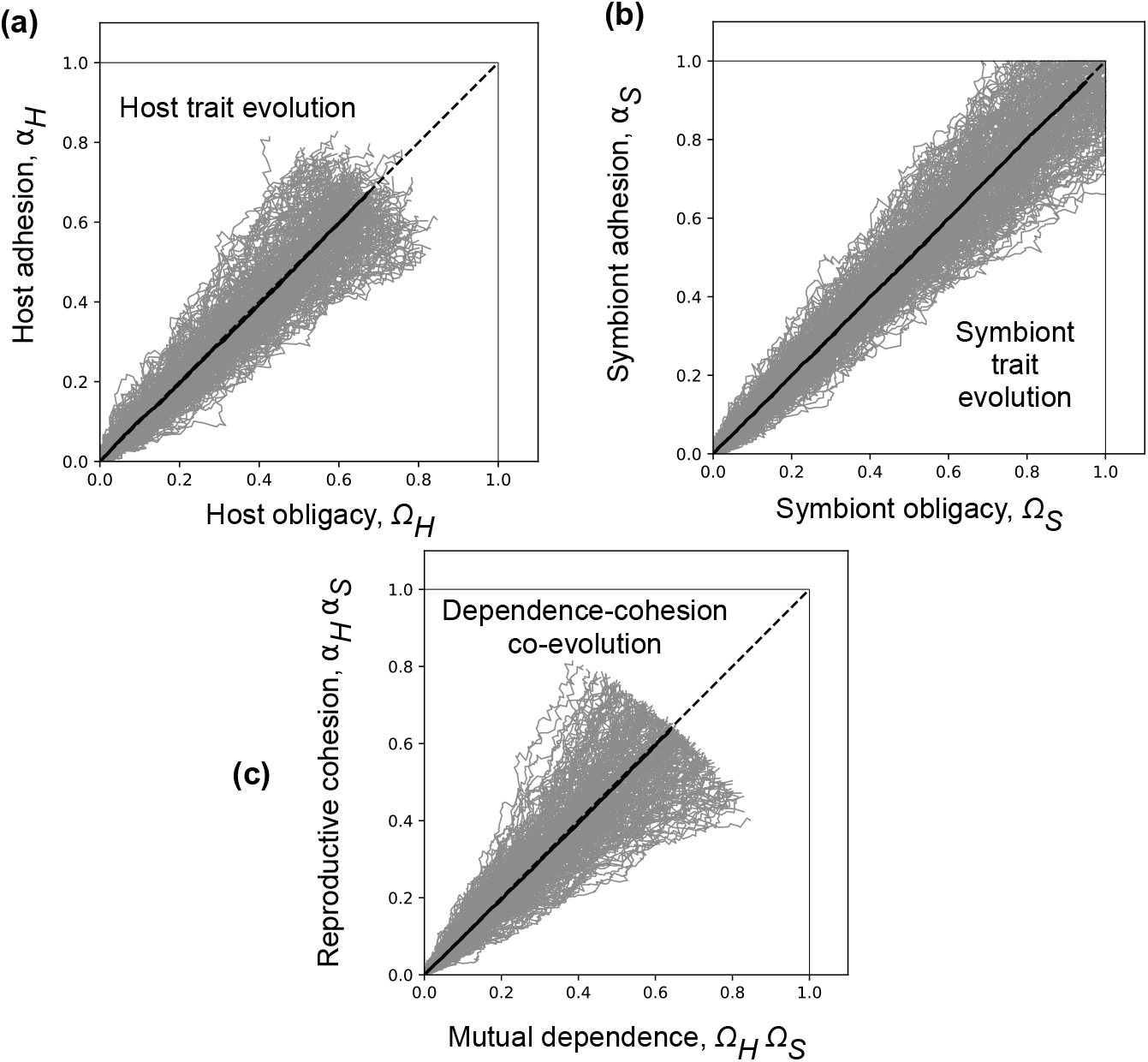
Independent growth rate depends on both obligacies, dissociation rate depends on both adhesions.

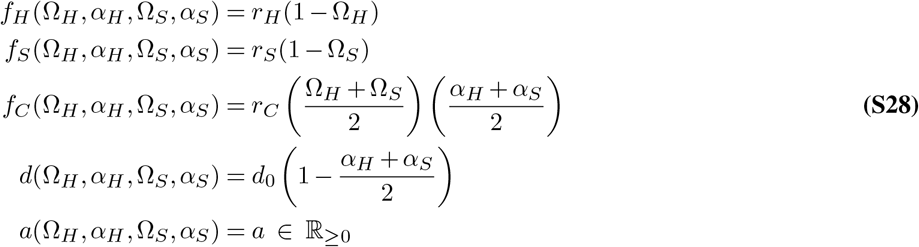

## Supplementary Note S5: Computation of the average trajectory in simulations with runs of different lengths

Across several figures of the main text and supplementary information, we present plots with several independent simulation runs of different time durations in grey, with a mean trajectory in solid black for interpretational convenience. When our exponential model of population dynamics is involved (Eq. S6), the lengths of these runs might be different. This is because they end when the feasibility boundary (eq. S9) is violated, and this can happen at different times because of the stochasticity of mutant traits. Here we briefly detail how we compute the average trajectory when runs can have different lengths.

Suppose there are *N*_*T*_ total trajectories (grey in the figures). Let *L*_*i*_ be the length of the *i*th trajectory, and suppose *L*_*max*_ = *max*_*i*_*L*_*i*_ is the duration (hereafter used interchangeably with length) of the longest trajectory, i.e. the trajectory that takes the most time. Denote trajectory *i* by a function *f*_*i*_(*t*) where *t ∈* {0, …, *L*_*i*_} (for example, from zero to the time it takes to reach the feasibility bound). We first artificially extend all trajectories (in time) such that they are all of length *L*_*max*_. In particular, we redefine for each *i*

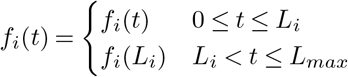

Recall that *t* cannot be larger than *L*_*max*_ for any trajectory by definition.

**Fig. S14.**
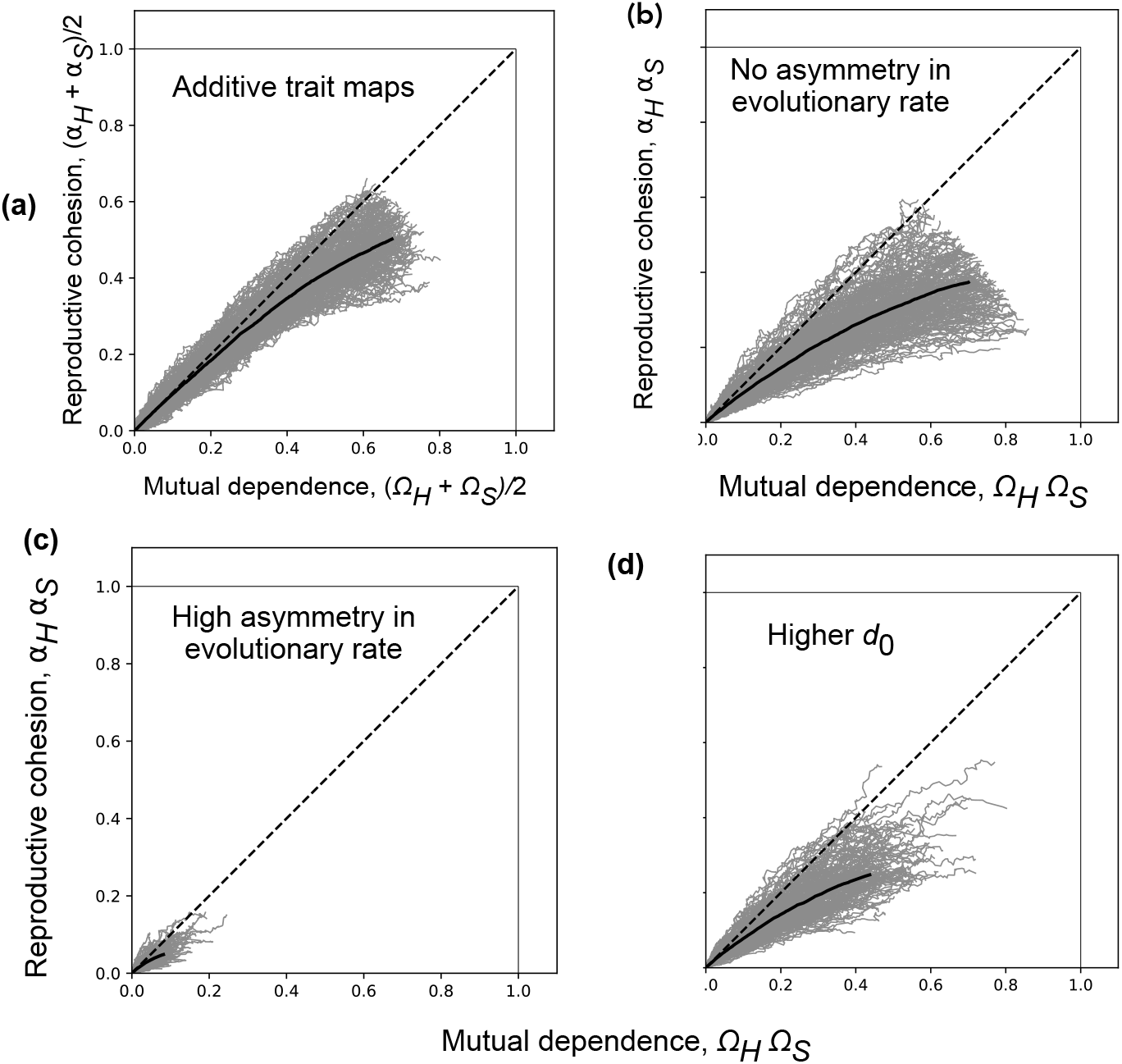
The bias towards mutual dependence is remarkably robust to model choices. We change important model choices to test the robustness of this result. None of the changes we have made affect its validity. **Base parameter values**. *fH* = 8, *fS* = 20, *rC* = 10, *KH* = 100, *KS* = 200, *a* = 0.1, *d*_0_ = 50. Changes are made to these values in the text that follows only when explicitly mentioned. **(a)** First, we change the map taking traits to ecological parameters into one where the traits have additive effects: the growth rate of the collective is proportional to the arithmetic mean of Ω*H* and Ω*S*, etc. The full map is presented in Eq. (E). **(b)** Here, we set *rH* = *rS* and *CH* = *CS* while keeping all other parameters fixed. **(c)** Now we set *rS* = 10*rH* and *CS* = 10*CH* to understand if a much higher asymmetry in the growth rates and carrying capacities affects evolutionary trajectories. **(c)** Lastly, we set *d*_0_ = 150 to understand if a much higher initial dissociation rate does something. It does not, really.

All trajectories are now of the same length, and the mean trajectory 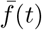 (in solid black) is calculated by

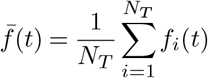

See the associated GitHub repository for the scripts that this is implemented in.

## Supplementary Note S6: A short note for reproducibility

In Figure 3 of the main text, we present the effects of varying the collective carrying capacity *K*_*C*_ and the growth rate scaling parameter *r*_*C*_. We conclude that *r*_*C*_ does not affect the results, whereas *K*_*C*_ both qualitatively and quantitatively affects the final evolutionary outcomes.

This outcome might sometimes be obscured during reproduction of our results: recall that, during our numerics, instead of the analytically true condition of merely *R*_0_ *>* 1 for the invasion of a mutant, we implement a cutoff of *R*_0_ *>* 1 + *ε* for a small positive value *ε* = 10^−7^. This is done to constrain the effects of provably (see Results section) artificial floating point errors which falsely lead to an *R*_0_ *>* 1 at the end of trajectories such as in Figure 2(b,e) of the main text.

However, when *r*_*C*_ is small, the eigenvalues giving rise to the values of *R*_0_ are also very small (smaller than the above 1 + *ε*). When comparing outcomes for different values of *K*_*C*_, these small *R*_0_ values are not accepted by the condition *R*_0_ *>* 1 + *ε* even though they lead to a systematic increase in the trait values (which can be checked by removing the *ϵ*).

We impose the criterion *R*_0_ *>* 1 + *ε* for expository clarity, but this criterion can be removed in the associated scripts – it would have the benefit of unambiguous demonstration of the fact that critical *K*_*C*_ values are *r*_*C*_-independent; but this is accompanied by the cost of artificial fluctuations of trait values even when *R*_0_ = 1 which might lead to confusion as well.

## Footnotes

1 The multiplicative form here is connected to how they map to ecological parameters - a multiplicative effect of the Ω_*i*_ on collective reproduction *f*_*C*_ motivates the interpretation of Ω_*H*_ Ω_*S*_ as the degree of mutual dependence as opposed to other potential measures such as Ω_*H*_ + Ω_*S*_.

